# Disrupted serial dependence suggests deficits in synaptic potentiation in anti-NMDAR encephalitis and schizophrenia

**DOI:** 10.1101/830471

**Authors:** H. Stein, J. Barbosa, M. Rosa-Justicia, L. Prades, A. Morató, A. Galan, H. Ariño, E. Martinez-Hernandez, J. Castro-Fornieles, J. Dalmau, A. Compte

## Abstract

We report markedly reduced working memory-related serial dependence with preserved memory accuracy in anti-NMDAR encephalitis and schizophrenia. We argue that NMDAR-related changes in cortical excitation, while quickly destabilizing persistent neural activity, cannot fully account for a reduction of memory-dependent biases. Rather, our modeling results support a disruption of a memory mechanism operating on a longer timescale, such as short-term potentiation.

---

The NMDA receptor (NMDAR) subserves memory mechanisms at several timescales, including sustained working memory delay activity ^1,2^ and different temporal components of synaptic potentiation ^3,4^. In addition, hypofunction of NMDARs is linked to psychiatric disease, in particular schizophrenia ^5^, and it possibly contributes to abnormal working memory function in patients with schizophrenia ^6,7^. Indeed, reduced prefrontal NMDAR density characterizes this disease ^8^. Yet, the specific neural alterations by which NMDAR hypofunction could lead to memory deficits in schizophrenia are still under debate ^6,7^. Here, we studied working memory function in healthy controls, patients with schizophrenia, and patients recovering from anti-NMDAR encephalitis (*Methods*, Supplementary Table 1). Anti-NMDAR encephalitis is characterized by an antibody-mediated reduction of NMDARs, accompanied by initial psychosis and long-lasting memory deficits ^9,10^, resembling clinical features of schizophrenia ^11^. Consequently, we expected working memory deficits in anti-NMDAR encephalitis to parallel those in schizophrenia. This correspondence allows linking alterations in working memory to the NMDAR in both patient groups.

We assessed memory alterations in a visuospatial delayed-response task (Fig. 1a) on two coexisting temporal scales: single-trial working memory accuracy as a proxy of active memory maintenance during short delays, and serial dependence of responses on previously memorized stimuli ^12,13^ (serial biases, Fig. 1b) as a read-out of passive information maintenance across trials. Neural correlates of this task have been identified in monkey prefrontal cortex ^14,15^, inspiring computational models that can capture key aspects of neural dynamics and behavior ^15–19^. The biophysical detail of these models permits to investigate how NMDAR hypofunction at different synaptic sites affects circuit dynamics and working memory. Candidate mechanisms are a disturbed balance between cortical excitation and inhibition (E/I balance), as it is observed in schizophrenia and in studies using NMDAR antagonists (e.g. ketamine) ^2,5,20,21^, and alterations in NMDAR-regulated short-term synaptic potentiation ^3,4,22^. In the modeling section of this study (Figs. 2,3), we systematically tested the potential of these candidate mechanisms for explaining experimentally observed memory alterations in schizophrenia and anti-NMDAR encephalitis.

**Fig. 1.**
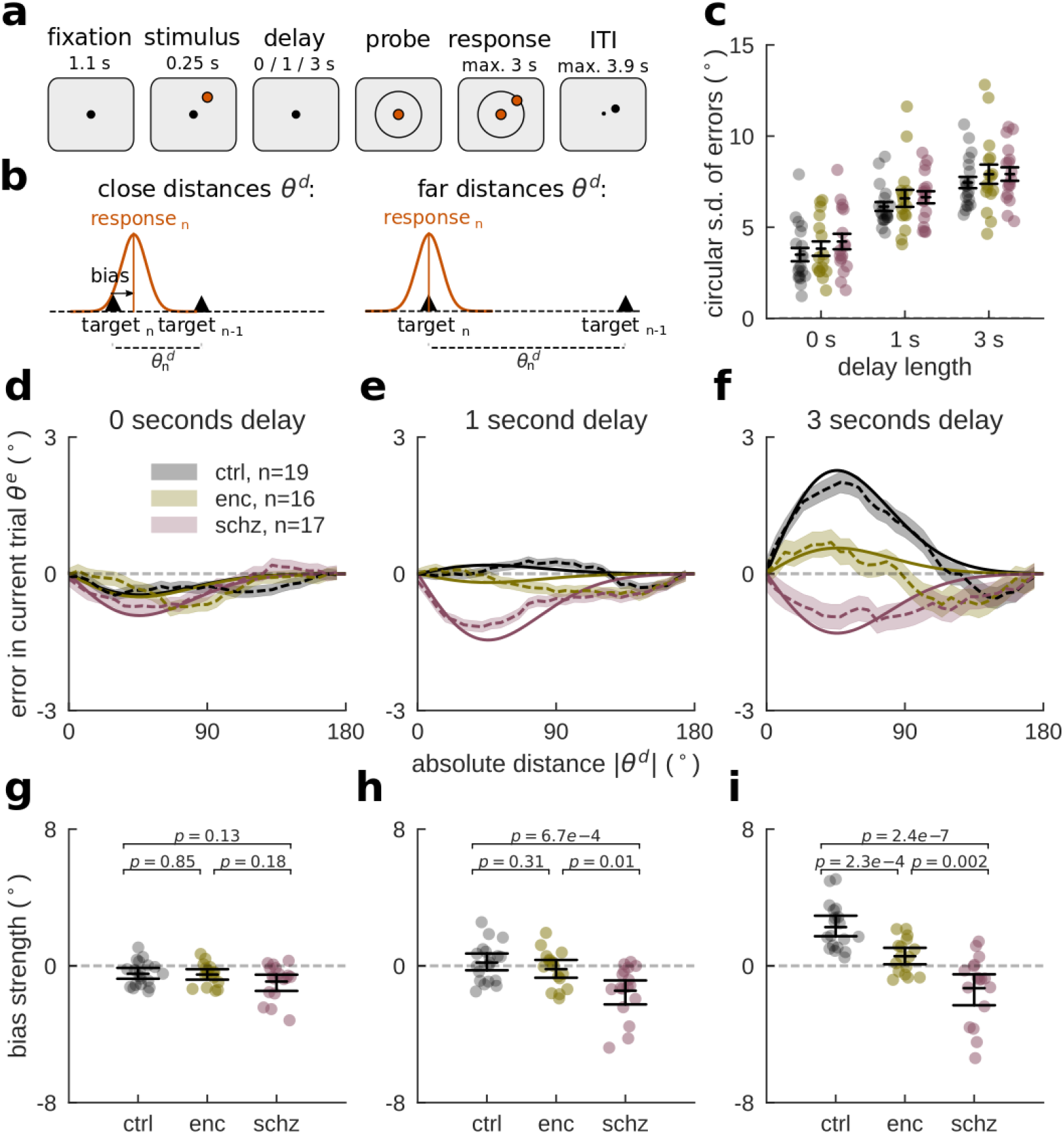
Reduced working memory-dependent serial dependence in anti-NMDAR encephalitis and schizophrenia. **a**, In each trial, subjects were to remember a stimulus that appeared for 0.25 s at a randomly chosen circular location with fixed distance from the center. Delay lengths varied randomly between trials (0, 1 or 3 s). Subjects made a mouse click to report the remembered location and started the next trial by moving the mouse back to the screen’s center during the inter-trial-interval (ITI). **b**, Serial dependence is measured as a systematic shift of responses towards previous target locations. Attractive effects depend on the distance *θ*^d^ between previous and current stimulus. **c,** Accuracy levels for each subject and delay were estimated as the circular standard deviation (SD) of bias-corrected error distributions (*Methods*). For longer delays, participants’ responses were less accurate (*delay*, F(2,147) = 77.04, p < 2e−16). There were no overall or delay-dependent group differences in accuracy (*group*, F(2,147) = 1.72, p = 0.18; *group ✕ delay*, F(4,147) = 0.07, p = .99). **d,e,f,** Serial dependence by group and delay length. Serial dependence is calculated as the ‘folded’ error *θ*^e^’ for different *θ*^d^ (dashed lines; *Methods*). Solid lines show linear model fits (*Methods*), omitting intercepts and negative values of *θ*^d^. Shading, ± s.e.m. *ctrl*: healthy controls, *schz*: schizophrenia, *enc*: anti-NMDAR encephalitis. **g,h,i** Individual (random coefficients; dots) and group estimates of serial bias strength (fixed effects; black error bars indicate mean and bootstrapped 95% C.I. of the mean) by delay. **g**, There was significant repulsive serial dependence in 0 s trials (*DoG(*θ*^d^)*, F(1,51.6) = 12.60, p = 0.0008) independently of group (*group ✕ DoG(*θ*^d^)*, F(2,51.6) = 0.45, p = 0.64). **h,** For 1 s trials, group differences in serial dependence emerged (*group ✕ DoG(*θ*^d^)*, F(2,48.2) = 6.57, p = 0.003) between ctrl and schz (t = 3.74, p = 6.7e−4) and enc and schz (t = 2.74, p = 0.01). **i**, After 3 s delay, both patient groups showed reduced biases compared to ctrl (*group ✕ DoG(*θ*^d^)*, F(2,49.3) = 15.68, p= 5.4e−6; ctrl vs enc, t = 4.14, p = 2.3e−4; ctrl vs schz, t = 6.43, p = 2.4e−7, and enc vs schz, t = 3.40, p = 0.002).

**Fig. 2.**
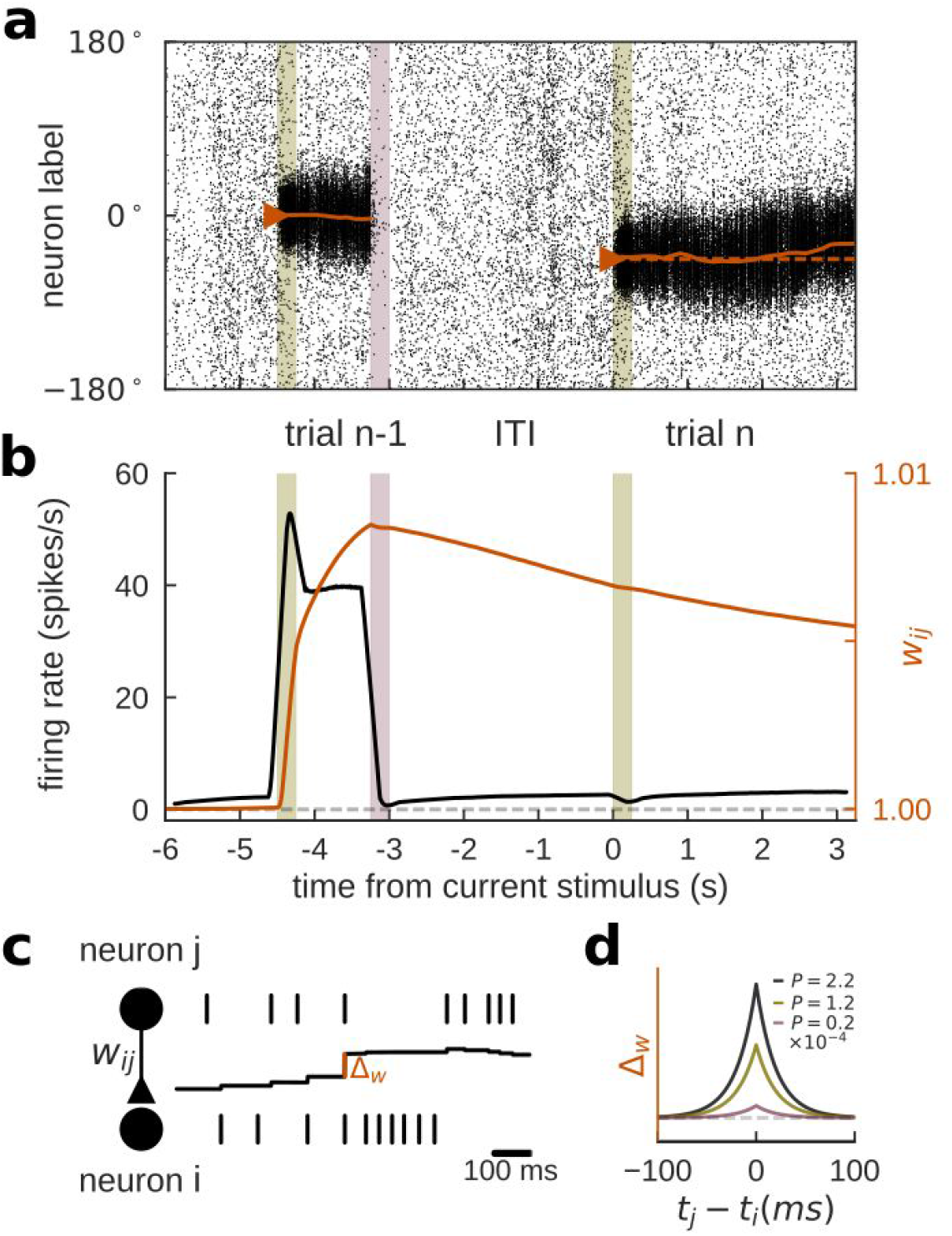
Ring attractor network with synaptic STP shows serial dependence. Simulations of two consecutive working memory trials (current trial *n*, previous trial *n-1*) in a spiking neural network model with bump-attractor dynamics (*Methods*). **a,** Spike times (x-axis) of excitatory neurons, ordered on y-axis by preferred angular location. Colored bars in **a,b** mark previous and current stimulus onset times (green) and previous response (red). The solid orange line shows the population vector decoded from firing rates (sliding windows of 250 ms). In trial *n*, the active memory representation got biased towards the memory representation in trial *n-1*. **b,** Firing rate (black) and potentiated weight trace *wij* for neuron at 0º (orange) averaged over 1,000 trials and 20 neurons centered around 0º. Spiking activity and synaptic strength increased during trial *n-1* delay and decreased after the response. At current stimulus onset, information about trial *n-1* remained only in the potentiated weight trace. To facilitate interpretation, we excluded trials for which any neuron participated in previous and current-trial delay activity (i.e., showed firing rates > 10 Hz after stimulus onset in trial *n*). **c,d,** Associativity and decay of modeled STP. The strength of each individual synapse is determined by *w_ij_* (**c,** middle black trace), which is potentiated at each spike by an amount *Δ* that depends on the relative spike times *t*_*j*_ and *t*_*i*_ of pre- and postsynaptic neurons, and on the potentiation factor *P* that is chosen to represent different strengths of STP (different colored lines in **d**; *Methods*, Eq. 14,15), and it is reduced by a fixed amount at each presynaptic spike, resulting in activity-dependent decay (Eq. 16).

**Fig. 3.**
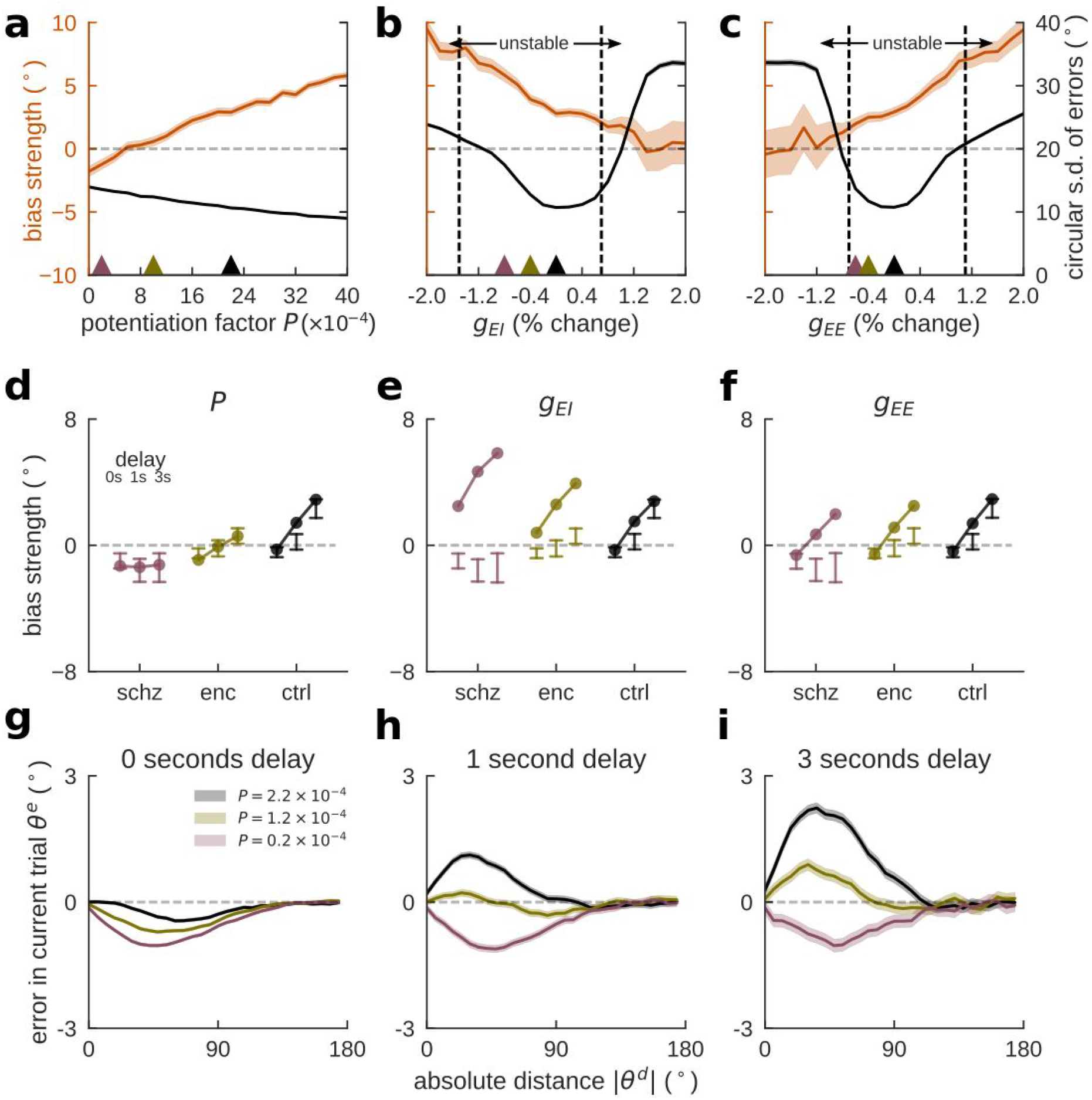
Altered STP simulates reduced serial dependence in spiking neural networks. **a,b,c,** Serial dependence (orange, bias coefficients from linear model, *Methods*) and accuracy (black, circular s.d. of errors) as functions of model parameters in 3 s delay trials (20,000 trials per parameter value). Vertical dashed lines indicate transition to ‘unstable’ network regimes for which more than 10% of trials were outliers (|*θ*^e^| > 57.3º, i.e. 1 radian). Shading, 95% C.I. **a,** Serial dependence decreased gradually when decreasing STP (potentiation factor *P*), while the network remained stable for all simulated values of *P*. Accuracy changed slightly as a function of STP. **b,** Cortical disinhibition via decreased *g_EI_* augmented serial bias while strongly affecting accuracy and stability, either due to instability of persistent activity (right, Supplementary Fig. 12b), or due to instability of baseline activity (left, Supplementary Fig. 12a). **c,** Lowering recurrent cortical excitation (*g_EE_*) led to the opposite pattern, decreasing biases. **d,e,f,** delay dependence of biases for each group, as defined by parameter values in a,b,c, (respectively colored triangles). For comparison, error bars indicate 95% CI for bias strength obtained from patients (reordered from Fig. 1g-i). **d,** Lowering STP strength reproduced the experimental data. In **e** and **f,** reduction of NMDAR conductances (*g_EI_* or *g_EE_*) did not reproduce group and delay dependencies of experimental biases. **g,h,i,** Serial dependence by delay length for different values of *P*, indicated by colored triangles in **a,** reproduced group- and delay-differences in bias observed in the data. Bias calculated as averaged ‘folded’ error *θ*^e^’ for binned absolute previous-current distances *θ*^d^. Shading, ± s.e.m. (20,000 trials per potentiation level *P*).

First, we sought to identify alterations in single-trial working memory accuracy, as an indication of a possible dysfunction of activity-based memory maintenance. Meta-analyses report mainly negative findings for delay-dependent accuracy impairments in schizophrenia and ketamine studies ^6,23^ (but see ref. ^24^). We calculated the circular standard deviation of bias-corrected response errors (*Methods*) as an inverse estimate of accuracy for each participant and delay. Correcting for biases as a systematic source of error allowed us to estimate memory accuracy independently of serial biases. For all groups, accuracy decreased equally with delay (Fig. 1c), indicating spared active working memory maintenance over short delays in encephalitis and schizophrenia.

Next, we tested whether NMDAR-related memory alterations could be observed at intermediate timescales by measuring serial dependence. Serial dependence is defined as a systematic shift of responses towards previously remembered, uncorrelated stimuli ^12^ (Fig. 1b), revealing that traces of recently processed stimuli persist in memory circuits and are integrated with new memories. Importantly, these attractive biases emerge over the trial’s memory delay, indicating a dependence on memory processes ^25,26^. In conditions without memory requirements, only small repulsive biases are present, possibly generated during perceptual processing ^25,26^. To assess NMDAR-related differences in serial dependence, we modeled single-trial errors 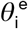 as a linear mixed model of delay length, group, and a non-linear basis function of the distance 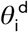 between consecutive stimuli ^12,26^ (derivative-of-Gaussian, DoG(*θ*^d^), *Methods*, Eq. 1; Supplementary Fig. 1).

In accordance with previous results ^25,26^, we found a dependence of attractive bias strength on memory delay (*delay ✕ DoG(*θ*^d^)*, (F(2,58) = 13.90, p = 1.2e−5). Moreover, biases differed between groups of participants (*group ✕ DoG(*θ*^d^)*, F(2,49) = 9.66, p = 0.0002), especially when comparing groups for different delay lengths (*group ✕ delay ✕ DoG(*θ*^d^)*, F(4,58) = 8.49, p = 1.8ee−5). Fig. 1d-f shows model fits and average bias curves for 0, 1 and 3 s delays (see Supplementary Fig. 2–4 for single-subject fits). Groupwise models (Eq. 2) allowed to assess the delay dependence of biases within each population (*delay ✕ DoG(*θ*^d^)*): For healthy controls, initially repulsive biases became gradually more attractive with delay length (F(2,16.9) = 26.91, p = 5.5e−6; Supplementary Fig. 5). Encephalitis patients showed a qualitatively similar, but reduced pattern (F(2,22.8) = 5.06, p = 0.015). In contrast, no attractive bias emerged over delay in patients with schizophrenia (F(2,18.1) = 1.61, p = 0.23). Rather, a repulsive bias dominated all delay lengths in this group (*DoG*(*θ*^*d*^), F(1,16.2) = 9.06, p = 0.008). Post-hoc tests and between-group comparisons are reported in Fig. 1g-i.

Serial dependence is known to fade with increasing inter-trial intervals (ITI) ^26^. We controlled for ITI length by including *ITI ✕ DoG*(*θ*^*d*^) as a covariate in our linear model (*Methods*, Eq. 4; Supplementary Fig. 6): For each additional second of ITI, serial bias decreased by 0.46±0.12º (mean±std). However, group differences in serial dependence remained unchanged. The timescale of serial dependence was further defined by how many past trials influenced the current response. We observed a much weaker delay-dependent bias towards the penultimate trial but there was no consistent evidence for group differences (Supplementary Fig. 7a-c).

We also controlled for potential effects of antipsychotic medication (*CPZ, Methods*). While CPZ correlated with bias strength in memory trials, delay-dependent biases still markedly differed between groups (Supplementary Fig. 8, upper left panels and caption). We did not find correlations between individuals’ bias estimates for 3 s delay trials and the severity of psychiatric symptoms for encephalitis or schizophrenia patients (Supplementary Fig. 8 and Supplementary Table 1). These between-subjects analyses were possibly underpowered, so we designed a within-subject longitudinal assessment for n=14 encephalitis patients that returned for a follow-up session after 3-12 months (mean 8.5 months). As expected, clinical symptoms improved in these patients (Supplementary Table 2) and we found that serial dependence normalized with the patients’ recovery (Eq. 7; Supplementary Fig. 9).

Together, our experimental results show no differences in single-trial memory maintenance, but a strong reduction of delay-dependent biases in anti-NMDAR encephalitis that ameliorates with patients’ recovery, and a complete absence of attractive biases in patients with schizophrenia. These findings are not explained by ITI length, general response correlations between trials (Supplementary Fig. 7d-f), or medication. Our conclusion is thus that alterations at the neural circuit level, related to NMDAR hypofunction, reduce serial dependence gradually, up to the point of completely disrupting attraction to previous stimuli. A prevailing idea associates NMDAR hypofunction in schizophrenia primarily to synapses onto GABAergic interneurons ^20^, while the role of NMDARs in working memory has been emphasized in synapses between pyramidal neurons ^1,2,18^. Alternatively, NMDARs could be involved in mechanisms directly associated with the generation of serial biases, such as short-term plasticity ^15,19,27^. To assess these mechanistic explanations comparatively, we simulated consecutive trials of a spatial working memory task in a spiking neural network model of the prefrontal cortex ^18^ (Fig. 2a). Prefrontal cortex not only holds working memory contents in an activity-based code ^14,17^, but also keeps long-lasting latent (possibly synaptic) memory traces that produce serial dependence ^15^.

We modeled a local prefrontal circuit, composed of neurons selective to the locations presented in the spatial working memory task. We used a network of excitatory and inhibitory neurons recurrently connected through AMPAR-, NMDAR- and GABA_A_R-mediated synaptic transmission in which persistent delay firing emerges from attractor dynamics (Fig. 2a, Supplementary Fig. 10; *Methods*). As proposed by previous studies ^15,19,27^ we modeled serial dependence as an effect of short-term plasticity that builds up at delay-active excitatory synapses and maintains information during the ITI in a subthreshold stimulus representation not reflected in firing rate selectivity (Fig. 2b, *Methods*). We modeled an associative mechanism of short-term potentiation (STP) that is NMDAR-dependent and upregulates glutamatergic efficacy, consistent with a long-lasting increase in the probability of presynaptic neurotransmitter release ^3,4^. As described, this efficacy increase undergoes activity-dependent decay ^3,4^ (Fig. 2c). In our simulations, stimulus-specific potentiated synaptic traces persisted through the ITI and attracted the next trial’s memory representation progressively over the course of the delay ^19,27^. To mimic memory-independent repulsive biases, current stimulus inputs were slightly shifted away from previous stimulus values by a fixed value ^27^ (*Methods*).

We assessed the effects of NMDAR dysfunction on serial dependence at three potential synaptic sites: based on the reported NMDAR-dependence of STP ^3,4^, NMDAR hypofunction would reduce the strength of STP at recurrent excitatory synapses and disrupt delay-dependent biases (hypothesis I: reduced STP). Also, we tested the explanatory potential of reduced NMDAR-mediated synaptic transmission. In particular, we tested cortical disinhibition ^24^, caused by diminished NMDAR efficacy at inhibitory interneurons (hypothesis II: reduced g_EI_), and the hypofunction of NMDARs at recurrent excitatory synapses, leading to diminished delay activity ^2^ (hypothesis III: reduced g_EE_). To assess each of these mechanisms, we independently varied STP strength, g_EI_ and g_EE_, and we read out “behavioral responses” after 0, 1 and 3 s from population activity in our network simulations (*Methods*). Then, we fitted a linear model to measure bias strength in each condition (Eq. 17, Supplementary Fig. 11). We sought to identify which mechanisms could independently reproduce the patterns of reduced and absent biases observed in patients, and their dependence on working memory delay (Fig. 1).

We found that both hypotheses I and III were qualitatively consistent with our experimental results: NMDAR hypofunction (whether reducing STP or g_EE_) reduced the strength of serial dependence (Fig. 3a,c, orange). In contrast, hypothesis II was discarded by our simulations: reducing g_EI_ increased serial dependence (Fig. 3b, orange), contrary to our experimental results, and quickly led to network disinhibition, causing previous-trial delay activity to spontaneously reemerge in the ITI (Supplementary Fig. 12). We noted that memory accuracy was slightly affected by all three manipulations (Fig. 3a-c), in contrast with our behavioral findings (Fig. 1b), but consistent with other studies with longer delays ^24^. Delay length and task complexity could be important factors to detect NMDAR-related differences in memory precision.

In addition, we found that hypotheses I and III could be disambiguated based on biases produced by the different models in 0, 1 and 3 s delays (Fig. 3d-f). Even for the lowest value of g_EE_ within the stable network regime, attractive biases increased with delay (Fig. 3f), contrary to our results for patients with schizophrenia (Fig. 1). In contrast, reduced STP reproduced equally strong repulsive biases for all delay lengths (Fig. 3d). Based on this modeling, we conclude that the disruption of STP, a mechanism operating on a longer timescale than activity-based memory maintenance, provides a plausible explanation for altered serial dependence as observed in schizophrenia and anti-NMDAR encephalitis.

In summary, we found a drastic reduction of working memory serial dependence in patients with anti-NMDAR encephalitis and schizophrenia, as compared to healthy controls. In contrast, we did not find memory maintenance deficits on timescales of a few seconds, suggesting that cognitive deficits in these patients ^7,10^ might be partly explained by the disruption of long-lasting, inactive memory traces, and a lacking integration of past and current memories. Our modeling results show that simple alterations in cortical excitation (hypotheses II and III), as proposed by current theories of NMDAR hypofunction in schizophrenia ^5,21,24^, cannot fully explain these behavioral findings. Instead, altered serial dependence is mechanistically accounted for by a disruption in slower dynamics, here specified as NMDAR-dependent associative STP (hypothesis I) that is triggered by sustained delay activity and influences memory representations in upcoming trials. Our results suggest that clinical reports of short-term memory alterations in schizophrenia and anti-NMDAR encephalitis could be understood in the light of reduced synaptic potentiation ^22^. This is consistent with *in vitro* studies, which have demonstrated the dependence of STP on specific subunit components of the NMDAR ^3,4^, and reduced STP in genetic mouse models of schizophrenia ^28^. Importantly, our modeling is not incompatible with altered cortical excitatory or inhibitory tone as a result of hypofunctional NMDARs. Rather, it states the necessity of assuming alterations in a mechanism operating on longer timescales, such as STP. For instance, diminished STP alongside symmetric effects on both E-E and E-I synapses could maintain the E/I balance and thus stable delay activity, while interrupting passive between-trial information maintenance.

Future studies should address the effects of pharmacological NMDAR blockade on serial dependence. These studies could unequivocally confirm the role of the NMDAR for trial-history effects in working memory, and at the same time allow to ask more specific questions: On the one hand, serial dependence effects under different NMDAR antagonists should vary according to how blocking specific NMDAR subunits modulates synaptic potentiation at different timescales ^3^. Our results cannot address subunit specificity because anti-NMDAR encephalitis (and possibly schizophrenia ^8^) is associated with hypofunction of the GluN1 subunit contained in all NMDARs ^11^. On the other hand, pharmacological studies in combination with neural recordings could reveal how trial-history representations are affected by the blockade of NMDARs ^15,29^. In rodents, long-term pharmacological experiments during behavior could be complemented with in vitro studies to assess STP directly. Finally, pharmacological studies would clarify if the alterations in serial dependence occur as a result of acute NMDAR hypofunction or whether they depend on compensatory changes in STP that arise after early, acute phases of cortical E/I imbalance in these diseases (e.g., as a long-term adjustment of the probability of presynaptic neurotransmitter release).

Our findings advance the conceptual understanding of working memory alterations in schizophrenia and anti-NMDAR encephalitis, as they demonstrate a selective disruption of information carryover between trials, reflected by a reduction of serial biases robustly found in neurotypical subjects ^13^. While we could not find correlations of reduced serial dependence with psychiatric scales (Supplementary Fig. 8), we showed that biases normalized with recovery from anti-NMDAR encephalitis (Supplementary Fig. 9). This suggests that serial biases reflect a clinically relevant dimension not captured in psychiatric scales. In this sense, it has been argued that serial dependence could facilitate information processing in temporally coherent real-world situations ^13^. Alternatively, serial biases could be the mere by-product of long-lasting cellular or synaptic mechanisms that support memory stabilization during working memory delays ^30^. Our study is in line with previous findings of reduced susceptibility to proactive interference in schizophrenia ^31^. However, while proactive interference is mainly discussed in the context of cognitive control, the limited complexity of our task restricts possible interpretations of reduced between-trial interference and supports the role of reduced residual memory traces. Moreover, thanks to our task’s well-studied single-neuron correlates ^14,15^ and biophysical models ^15,17,18^ and the comparison with anti-NMDAR encephalitis patients, we provide a specific mechanistic model of synaptic deficits leading to reduced previous-trial interference in schizophrenia.

Interestingly, a reduction in serial dependence has recently been reported for patients with autism ^32^, a disease also associated with NMDAR hypofunction ^33^ and alterations in synaptic potentiation ^22^. Further, as for autism, our findings of reduced serial dependence are compatible with normative accounts of information processing in schizophrenia. Classic theories and recent studies have reported an underweighting of past context, or in Bayesian terms, learned priors, and an overweighting of incoming perceptual information in patients with schizophrenia ^34–36^ and NMDAR hypofunction ^37^. Long-lived traces of past stimuli could serve as Bayesian priors to perception and memory, and a disruption of STP might be regarded as a biological implementation of a reduced usage of priors in schizophrenia and anti-NMDAR encephalitis.

## Acknowledgments

We acknowledge support from Institute Carlos III, Spain (Ref: PIE 16/00014), CELLEX Foundation, Safra Foundation, CERCA Programme/Generalitat de Catalunya, Generalitat de Catalunya (AGAUR 2014SGR1265, 2017SGR01565), “la Caixa” Foundation (ID 100010434), under the agreement LCF/PR/HR17/52150001, and by the Spanish Ministry of Science, Competitiveness and Universities co-funded by the European Regional Development Fund (Refs: BFU 2015-65318-R, RTI2018-094190-B-I00). HS was supported by the “la Caixa” Banking Foundation (Ref: LCF/BQ/IN17/11620008), and the European Union’s Horizon 2020 Marie Skłodowska-Curie grant (Ref: 713673). We thank the Barcelona Supercomputing Center (BSC) for providing computing resources. This work was developed at the buildings *Centro Esther Koplowitz*, *CELLEX*, and *Hospital Clinic*, Barcelona. We thank Thaís Armangue, Domingo Escudero, Amaia Muñoz-Lopetegi and Gisela Sugranyes for assistance in recruiting patients. Jaime de la Rocha, Daniel Linares for discussions, Ainhoa Hermoso-Mendizabal for comments on the manuscript, and Diego Lozano-Soldevilla for assistance during the development of the task.

## Data and Code availability

Behavioral data and custom code generated or analysed for this work are available upon request.

## Author Contributions

HS, JB and AC designed behavioral and computational aspects of the study. JC, JD, and AC designed clinical aspects of the study. HS performed analyses of human behavior and computer simulations. HS, JB, and AC developed the computational model. HS, AM, LP and AG performed human experiments. MR and LP performed psychiatric testing. JC, MR, HA, EM recruited participants for the study. HS and AC wrote the manuscript. HS, JB, JD and AC discussed the results and edited the manuscript. All authors reviewed the manuscript for intellectual content.

## Methods

### Experimental Procedures

#### Sample

We included n=16 patients with anti-NMDAR encephalitis (*enc*), n=17 patients with schizophrenia or schizoaffective disorder (n=12 and n=5, respectively; *schz*), and n=19 neurologically and psychiatrically healthy control participants (*ctrl*), all with normal or corrected vision. Psychiatric diagnoses (or the absence thereof for controls) were confirmed using the Structured Clinical Interview for DSM IV (SCID-I) ^38^. Patients diagnosed with anti-NMDAR encephalitis were recruited from different centers (n=14 in Spain, n=1 in Germany and n=1 in the United Kingdom) at the moment of hospital discharge and completed the experiment around 5.5 months after disease onset (median, interquartile range i.q.r. = 3.5 months). All patients fulfilled clinical diagnostic criteria of anti-NMDAR encephalitis with confirmation of CSF IgG antibodies against the GluN1 subunit of the NMDAR ^39^. All subjects were tested in our laboratory for antibodies against NMDAR in serum as previously described ^40^ and all healthy controls and patients with schizophrenia were seronegative. Anti-NMDAR encephalitis is known to have a prolonged process of recovery after the acute stage of the disease ^41^, and patients included in this study still suffered from cognitive deficits (data not shown) as has been previously described in cohorts with long follow-up ^10^ but were able to participate in the testing procedure. Controls and patients with schizophrenia were recruited from the Barcelona area and from Hospital Clínic (Barcelona, Spain), respectively. Patients with schizophrenia were tested 35.0 months after diagnosis (median, i.q.r.=51.0 months) and were clinically stable at the time of testing. All participants (and, in the case of minors of age, their legal guardians) provided written informed consent and were monetarily compensated for their time and travel expenses, as reviewed and approved by the Research Ethics Committee of Hospital Clínic. All subjects were assessed for psychiatric symptoms and functionality through a battery of standard tests including the Spanish versions of the Positive and Negative Syndrome Scale (PANSS) ^42^, the Young Mania Rating Scale (YMRS) ^43^, the Hamilton Depression Rating Scale (HAM-D) ^44^ and the Global Assessment of Functioning Scale (GAF) ^45^. Finally, the dose of antipsychotic medication at the moment of testing was estimated as chlorpromazine equivalent (CPZ) ^46^. For a demographic and clinical overview of the populations, please refer to Supplementary Table 1.

#### Task protocol and behavioral testing

Participants completed two 1.5 h sessions performing a visuospatial working memory task described in Fig. 1a. In each session, participants were asked to complete 12 blocks of 48 trials. However, some participants did not complete all blocks (on average, participants completed 1114.1±134.4 trials (mean±std, ctrl), 1086.0±189.9 trials (enc), and 1030.6±192.8 trials (schz).

Each trial began with the presentation of a central black *fixation* square on a grey background (0.5 × 0.5 cm) for 1.1 sec. A single colored circle (*stimulus*, diameter 1.4 cm, 1 out of 6 randomly chosen colors with equal luminance) was then presented during 0.25 s at one of 360 randomly chosen angular locations at a fixed radius of 4.5 cm from the center. The stimulus was followed by a randomly chosen *delay* of 0 (16.67% of trials), 1 (66.67% of trials), or 3 s (16.67% of trials) in which only the fixation dot remained visible (except for 0 second trials, where the stimulus remained visible until the participant started to move the cursor). When the fixation dot changed to the stimulus’ color (*probe*), participants were asked to respond by making a mouse click at the remembered location (*response*). A white circle indicated the stimulus’ radial distance, so participants only had to remember the angular position. After the response, the cursor had to be moved back to the fixation dot to start a new trial (*ITI*). Participants were instructed to maintain fixation during the fixation period, stimulus presentation, and memory delay and were free to move their eyes during response and when returning the cursor to the fixation dot.

### Data analysis

#### Error and serial dependence analysis

Response errors 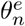 in trial *n* were measured as the angular distance between response and target. To exclude errors due to guessing or motor imprecision, we only analyzed responses within an angular distance of 1 radian and a radial distance of 2.25 cm from the stimulus. Further, we excluded trials in which the time of response initiation exceeded 3 s, and trials for which the time between the previous trial’s response probe and the current trial’s stimulus presentation exceeded 5 s. In total, 2.6±4.2% (mean±std, ctrl), 4.8±6.9% (enc) and 7.5±9.6% (schz) of trials per participant were rejected (but only 0.1±0.2% (ctrl), 0.4±0.5% (enc) and 0.6±0.7% (schz) of trials were excluded due to angular response errors).

We then measured serial dependence as the error in the current trial as a function of the circular distance between the previous and the current trial’s target location. Fig. 1c,d,e depict ‘folded’ serial dependence: We multiplied trial-wise errors 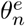 by the sign of the previous-current distance, 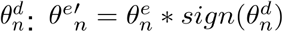, and then binned data based on absolute values 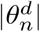. Errors 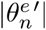 were then averaged for each 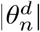 in sliding windows with size 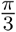 in steps of 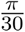. Positive mean folded errors should be interpreted as attraction towards the previous stimulus and negative mean folded errors as repulsion away from the previous location. For visualization, all values were transformed from radians to angular degrees.

#### Linear (mixed) models

We modeled signed errors 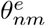 in trial *n* and subject *m* using a linear mixed model that included the factors *group* (ctrl, enc or schz), *delay* (0, 1, or 3 s) and a nonlinear function of previous-current stimulus distance 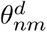, 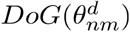, which has been used for modeling serial dependence ^12,26^. It is the normalized first derivative of a Gaussian with fixed location hyperparameter *μ* = 0. The variance hyperparameter *σ* was determined using cross-validation as explained below (see also Supplementary Fig. 1). Our main model is:

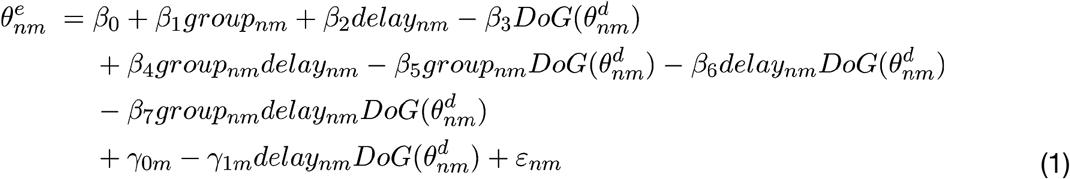

*β* coefficients estimate fixed, and *γ* coefficients random effects. Bias strength for a certain condition can then be read out as the sum of coefficients of all terms containing 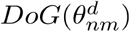, and the dependence of bias strength on other variables is assessed by evaluating the significance of interaction terms containing 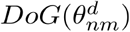 and the relevant variable. To measure response accuracy, bias-corrected response errors were defined as model residuals *ε*_*nm*_ from Eq. 1. For each subject and delay, response accuracy was then measured as the circular s.d. of *ε*_*nm*_.

Group- (Supplementary Fig. 5) and delay-wise (Fig. 1g-i) models were defined as:

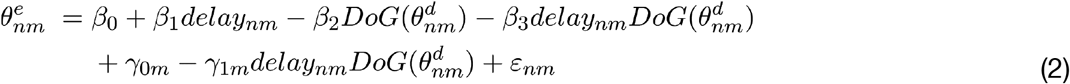

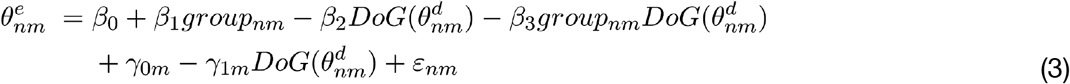

The effect of covariates *ITI* length (Eq. 4) and *CPZ* equivalent (Eq. 5) were assessed as:

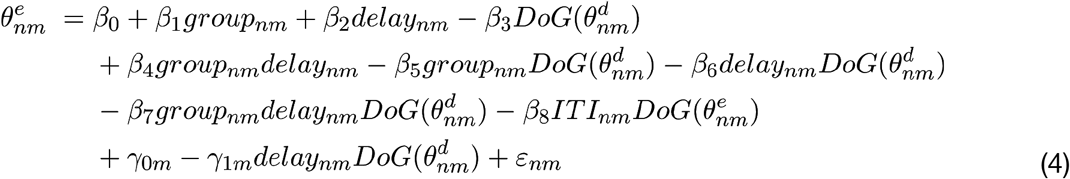

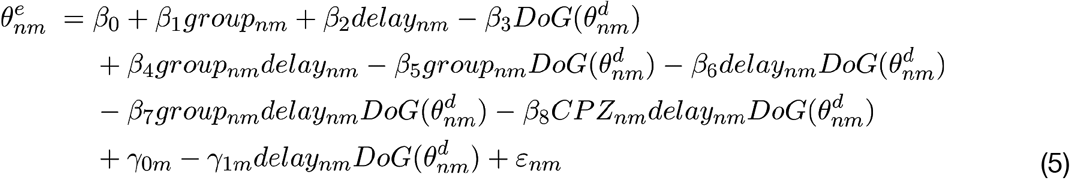

Biases towards stimuli in trial *n-2* were measured by including distances to the penultimate stimulus, 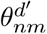 :

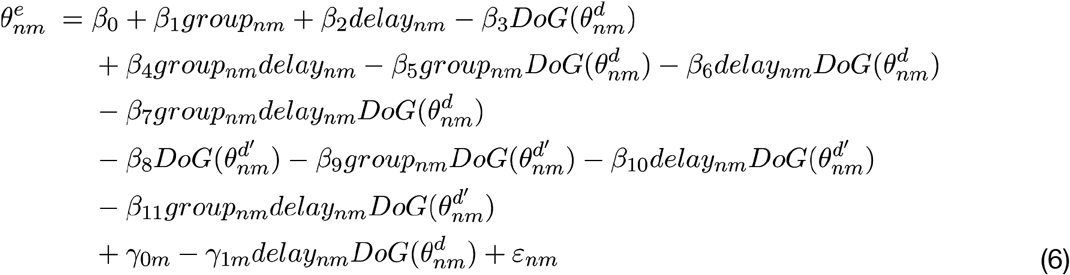

Baseline- and follow-up sessions in encephalitis patients and controls were compared by:

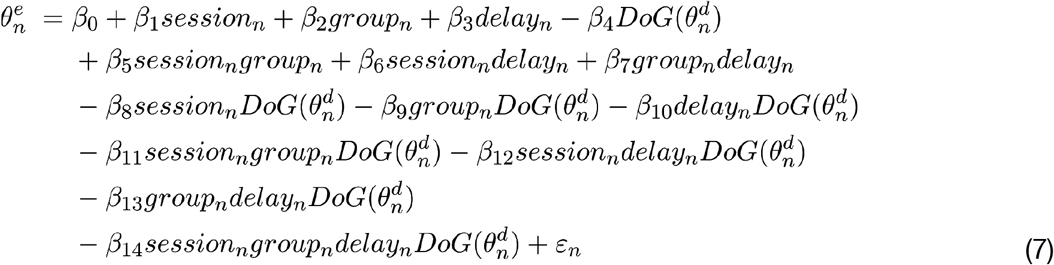

For extended models in Eq. 4–7, we compared nested models via Wald Tests to determine the optimal model complexity. All mixed models were fitted, compared and statistically tested using the R packages *lme4* ^47^ and *lmerTest* ^48^ through *rpy2*.

#### Cross-validation and hyperparameter fitting

To determine hyperparameter σ used in Eq. 1–7, we fitted errors 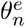 in trial *n* as a linear model including factors group, delay, and 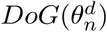 as described in Eq. 1, but excluding random effects:

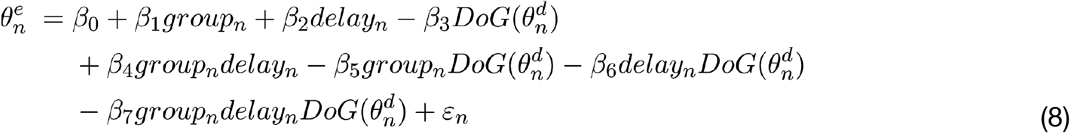

while setting Gaussian hyperparameters and *μ* = 0 and *σ* ∈ [0.2, 1.8] (in radians). For each value of variance *σ*, we used a stratified cross-validation procedure, fitting the model to 67% of the trials from each subject and testing the prediction in the left-out 33% of trials. Performance for each *σ* was evaluated using the mean squared error (MSE) of predictions from 1,000 cross-validation repetitions. *σ* was chosen as to minimize the model’s MSE, yielding *σ* = 0.8 (Supplementary Fig. 1).

#### Model selection

To test whether a model with repulsive biases at high distances 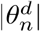 fitted our data more parsimoniously, we compared cross-validation MSE for models with first- and third-derivative-of-Gaussian basis functions (Supplementary Fig. 1). We repeated the hyperparameter fitting procedure described above for the third-derivative-of-Gaussian model using hyperparameters *μ* = 0 and *σ* ∈ [0.6, 2.0] rad. As the first-derivative-of-Gaussian model produced smaller MSE in the cross-validation procedure, we discarded the third-derivative-of-Gaussian model. Thus, all model results reported in this manuscript correspond to the first-derivative-of-Gaussian model.

### Neural network simulations

#### Network architecture and dynamics

We simulated consecutive pairs of trials in a spiking neural network model of prefrontal cortex implemented in Brian2 ^49^. *N*_*E*_ = 1024 excitatory and *N*_*I*_ = 256 inhibitory leaky integrate-and-fire neurons were connected all-to-all via synapses governed by NMDAR-, AMPAR-, and GABA_A_R-dynamics, as described in ref. (^18^).

The dynamics of the membrane voltage of excitatory neurons *V*_*i*_(*i* = 1‥*N*_*E*_) were given by:

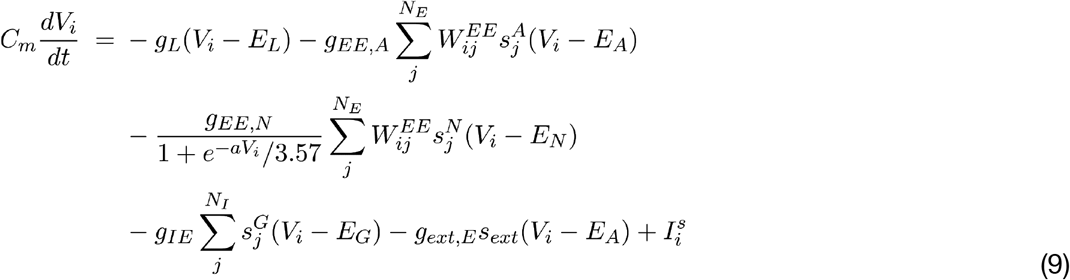

with membrane capacitance *C*_*m*_ = 0.5 nF, leak conductance *g*_*L*_ = 25nS,, leak reversal potential, *E*_*A*_ = 0 mV, AMPAR, GABA_A_R and NMDAR reversal potentials *E*_*A*_ = 0 mV, *E*_*G*_ = −70 mV, *E*_*N*_ = 0 mV, unitary, conductances *g*_*ext*_ = 3.1 nS, *g_IE_* = 2.672 nS, *g_EE,N_* = 0.56 nS, *g_EE,A_* = 502 nS, the NMDAR magnesium block parameter *a* = 0.062 mV^−1^. In simulations of reduced NMDAR conductance, parameters or respectively *g_EI,N_* were modulated as indicated in Fig. 3b,c,e,f and Supplementary Fig. 12.

The membrane voltage of inhibitory neurons followed:

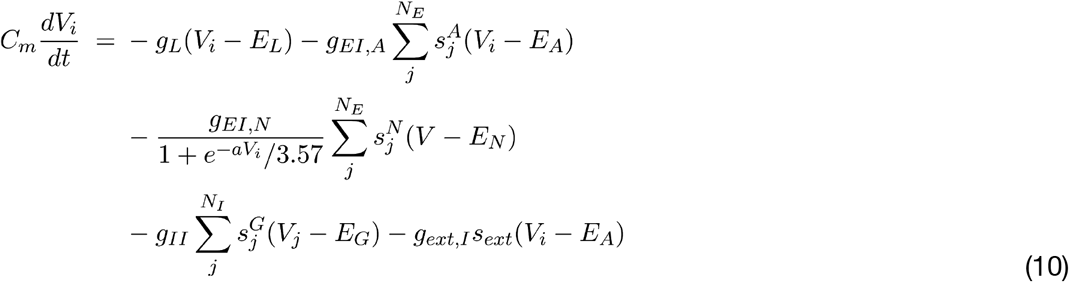

with *C*_*m*_ = 0.2 nF, *g_L_* = 20 nS, *g_ext,I_* = 2.38 nS, *g_II_* = 2.048 nS, *g_EI,A_* = 0.384 nS and *g_EI,N_* = 0.424 nS.

The kinetics of synaptic variables 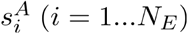, 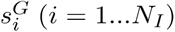 and *s*_*ext*_ were determined by

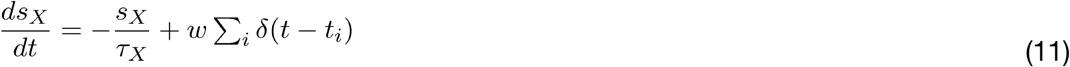

with *τ*_*A*_ = 2 ms, *τ*_*G*_ = 10 ms, *τ*_*ext*_ = 2 ms, and the summation runnin so that at each spike time the synaptic variable increased by a step of magnitude *w*, which was generally set to 1 except for synapses undergoing synaptic potentiation (see below). For *s*_*ext*_, spike times were generated as a Poisson spike train of rate 1800 sp/s (simulating inputs from 1,000 external Poisson neurons firing at 1.8 sp/s each).

The slower and saturating NMDAR synaptic variables 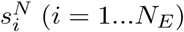 followed the coupled equations:

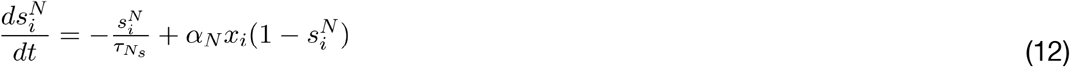

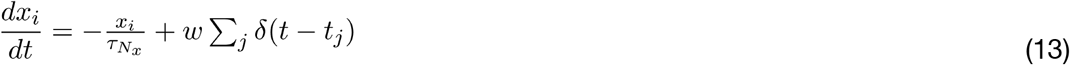

with 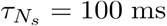, 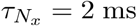, and *α*_*N*_ = 0.5 kHz.

The strength of recurrent excitatory synapses was modulated depending on the distance in preferred location of presynaptic and postsynaptic excitatory neurons: 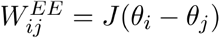, where *J* is a Gaussian function (centered at *μ* = 0 with s.d. *σ* = 14.4 deg) plus a constant, tuned so that Σ_*j*_ *J*(*θ*_*i*_ − *θ*_*j*_) = *N*_*E*_ and *J*(0) = 1.63. As a result, neurons with similar preferred locations had 1.63 stronger weights than the average weight (Supplementary Fig. 10 for network scheme and weight profiles).

#### STP rule

For connections between excitatory neurons, the spike-triggered step in AMPAR and NMDAR synaptic variables *w* could vary individually for each specific connection: *w*_*ij*_ characterized the step at the synapse from neuron *j* onto neuron *i*. Upon synchronized pre- and postsynaptic spiking, *w*_*ij*_ was slightly enhanced by an amount Δ_*w*_ that depended on the relative spike times of neuron *j* and *i* (Fig. 2c) to simulate an increase in probability of glutamate release ^50^:

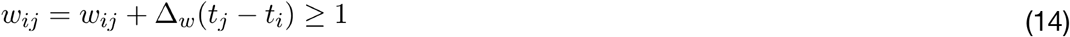

The associative nature of this rule was determined by a potentiation function that required synchronization within a specific temporal window (Fig. 2d):

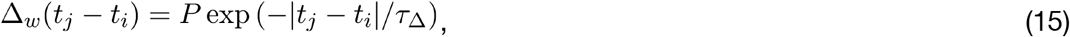

with potentiation factor *P* = 0.00022 and *τ*_Δ_ = 20 ms. Changes were sustained (did not decay with time), but synapses depotentiated based on presynaptic activity ^3^: at each presynaptic spike

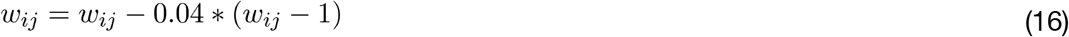

#### Trial structure and simulations

We simulated 20,000 pairs of consecutive trials with independent randomized stimulus locations. Network inputs 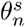 in trial *n* with stimulus *s* were slightly transformed to mimic a repulsive “baseline” bias away from previous stimulus locations, resulting from sensory aftereffects produced in lower-level cortical areas^26^: 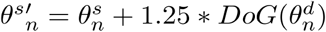, where 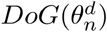 is the first-derivative-of-Gaussian function with **σ** = 0.8 rad and *μ* = 0, and 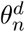 is the distance between previous and current stimulus.

Simulations started with a stimulus presentation at 0º (trial *n−1*) for 0.25 s. After the input was removed, a delay of 1 s followed. A negative input to the whole network during 0.25 s simulated the response and removed stimulus-associated neural activity. After an ITI of 3 s, a second stimulus (trial *n*) was delivered at a random location for 0.25 s. The second delay duration was 3 s. To obtain behavioral readouts from the network, we counted each neuron’s spikes during three time windows of 0.25 ms: 0-0.25 s after stimulus offset (0 s delay condition), 0.75-1 s (1 s delay), and 2.75-3 s after stimulus offset (3 s delay). The behavioral response was determined as the angular direction of the population vector of spike counts.

#### Network behavioral analysis

We first calculated the percentage of outlier responses and excluded outlier trials from the network’s population vector responses (response error > 1 radian). Circular standard deviations and serial dependence were then calculated from the network’s population vector responses analogous to human error analyses. In Fig. 3a-f, bias strength was measured as the sum of bias term coefficients in the linear model

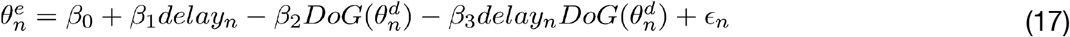

that fitted errors 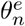 in trial *n* from each parameter manipulation (*P*, *g*_*EE*_, and *g*_*EE*_) separately as a function of delay and 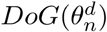 with and rad.

#### Cross-validation and hyperparameter fitting

The value of hyperparameter **σ** was determined in a cross-validation procedure for the “baseline” condition with *P* = 0.00022, *g_EE_* = 0.056 nS, and *g_EI_* = 0.424 nS, for values **σ** ∈ [0.2, 1.8] (in radians). For each value of **σ**, we fitted the model described in Eq. 14 to a 67% of trials and testing the prediction in the left-out 33% of trials. Performance for each was evaluated using the mean squared error (MSE) of predictions from 10 cross-validation repetitions. **σ** was chosen to minimize the model’s MSE, yielding **σ** = 0.60 rad (Supplementary Fig. 11).

## Supplementary Figures

**Supplementary Fig. 1.**
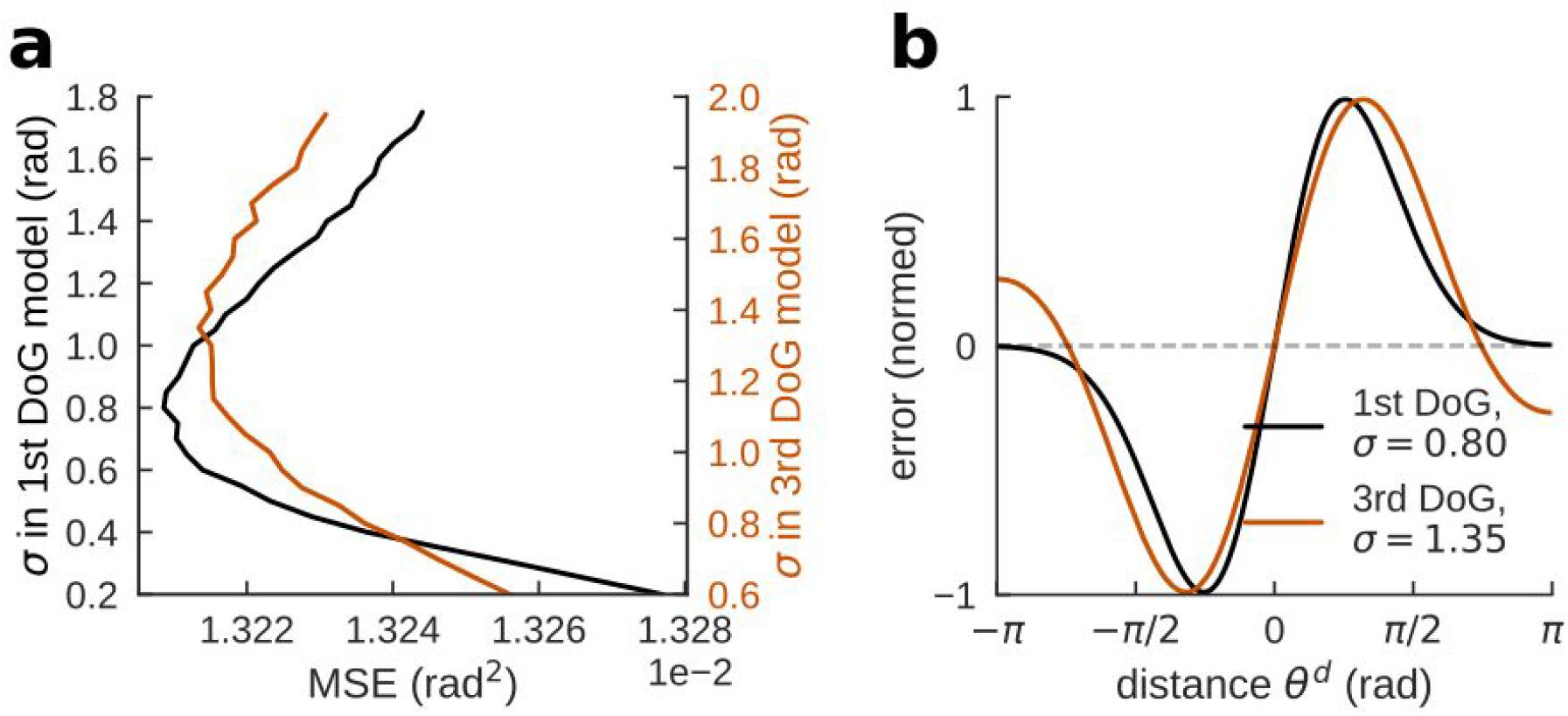
Hyperparameter cross-validation and model selection. **a,** Mean squared error for stratified hyperparameter optimization using cross-validation (1.000 repetitions, training set size = .33 from each subject) for first- (black) and third- (orange) derivative-of-Gaussian fits. Hyperparameters are different values of variance *σ* of the underlying Gaussian with location hyperparameter *μ* = 0. MSE: mean squared error. **b,** Shape of first- and third-derivative-of-Gaussian fits with optimal hyperparameter *σ* and *μ* = 0. The cross-validation procedure used for model selection was carried out based on a model with a minimal set of variables (group, delay, and DoG(*θ*^d^)), excluding random effects (*Methods*, Eq. 8). Note that signed previous-current distances in radians were used in the linear model.

**Supplementary Fig. 2.**
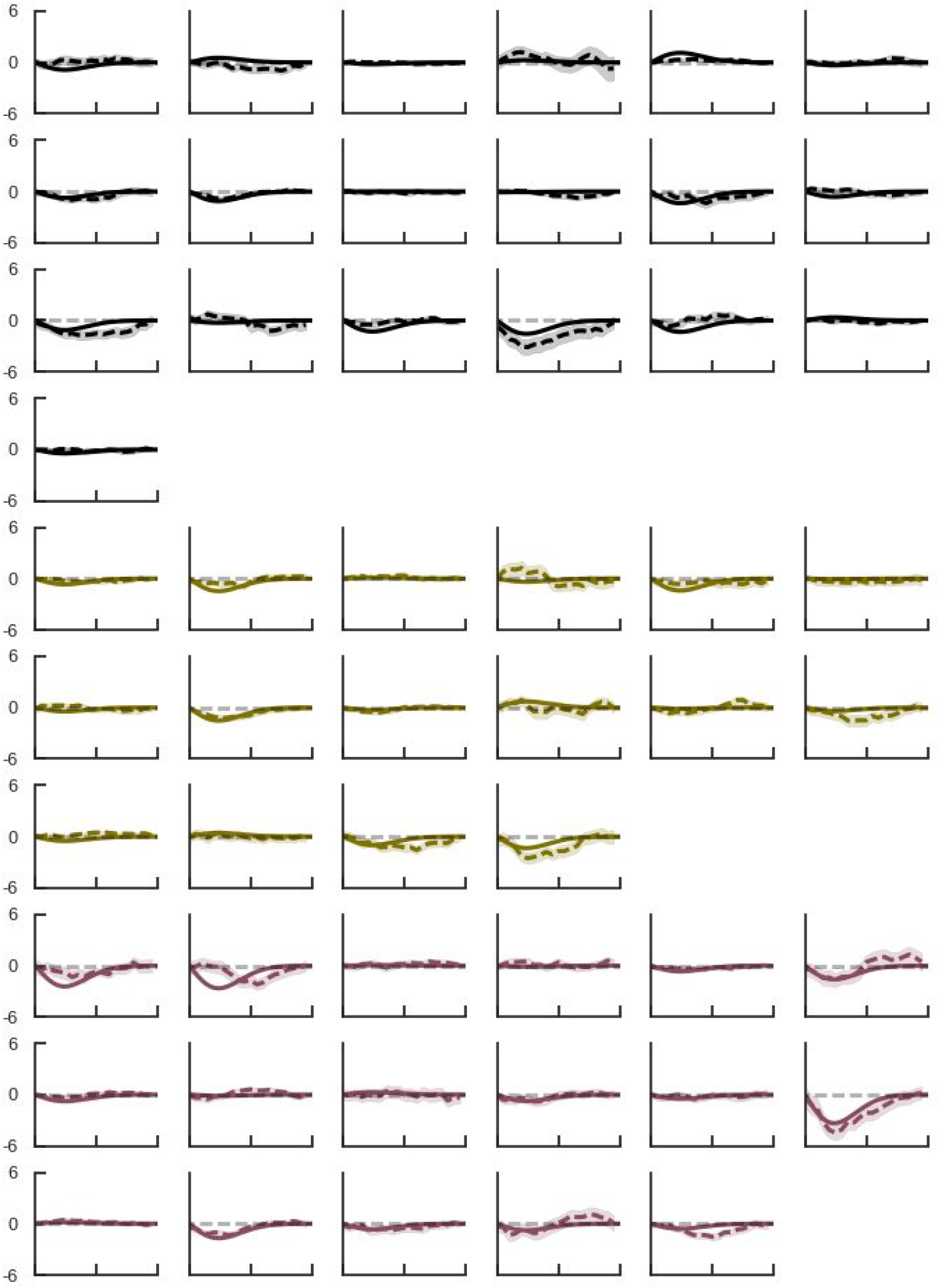
0 sec delay single subject bias and linear mixed model fit. Serial dependence is calculated for each subject as the ‘folded’ error *θ*^e^’ (in degrees, y-axis) for different previous-current distances *θ*^d^ (x-axis, spanning absolute values of 0°-180°) (dashed line; *Methods*). Shading, ± s.e.m. Solid lines show linear model fits (*Methods*, Eq. 1), omitting intercepts and negative values of *θ*^d^ for visualization. Black curves (row 1-4), *ctrl*, green curves (row 5-7), *enc*, purple curves (8-10), *schz*.

**Supplementary Fig. 3.**
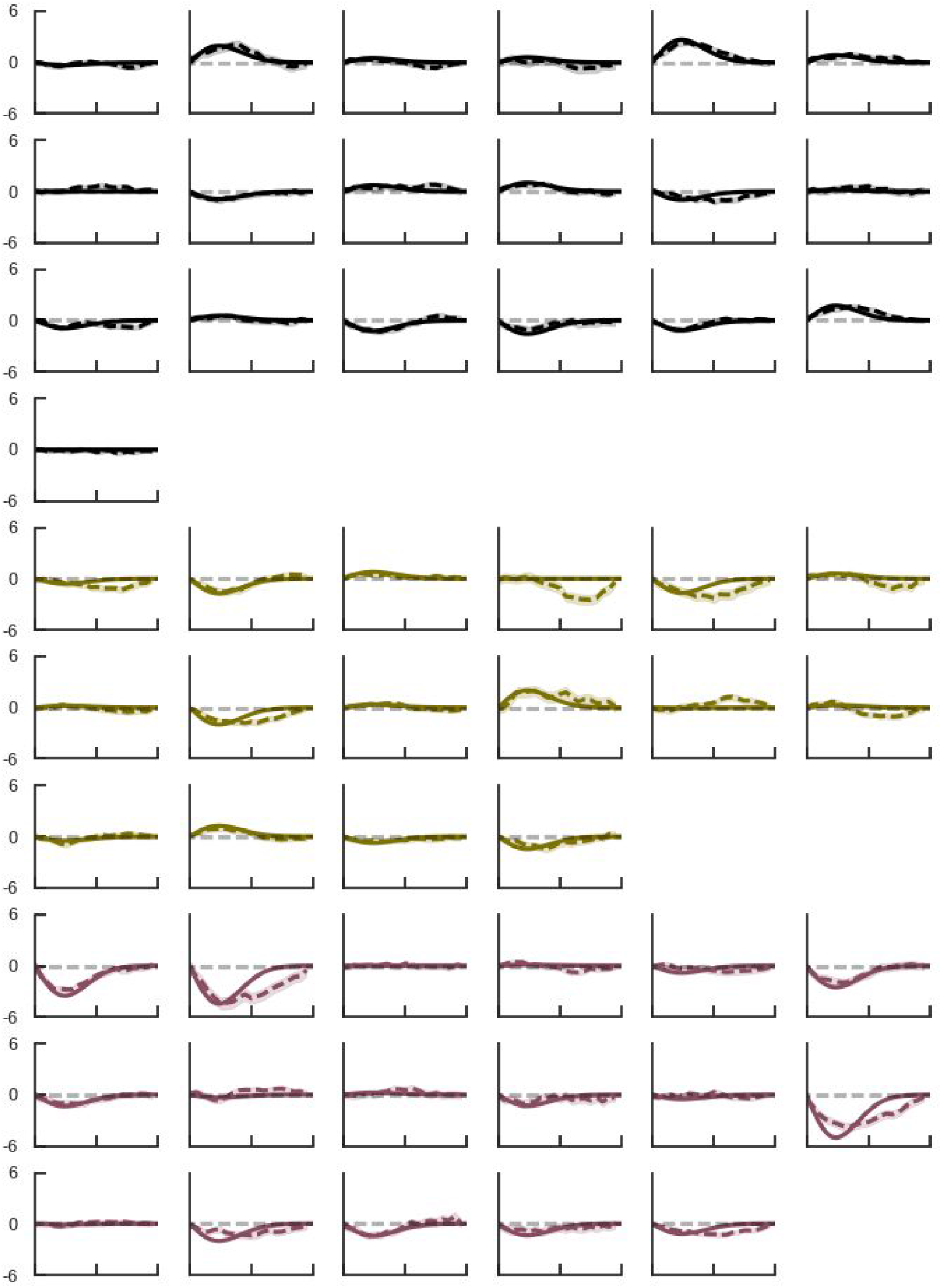
1 sec delay single subject bias and linear mixed model fit. Serial dependence is calculated for each subject as the ‘folded’ error *θ*^e^’ (in degrees, y-axis) for different previous-current distances *θ*^d^ (x-axis, spanning absolute values of 0°-180°) (dashed line; *Methods*). Shading, ± s.e.m. Solid lines show linear model fits (*Methods*, Eq. 1), omitting intercepts and negative values of *θ*^d^ for visualization. Black curves (row 1-4), *ctrl*, green curves (row 5-7), *enc*, purple curves (8-10), *schz*.

**Supplementary Fig. 4.**
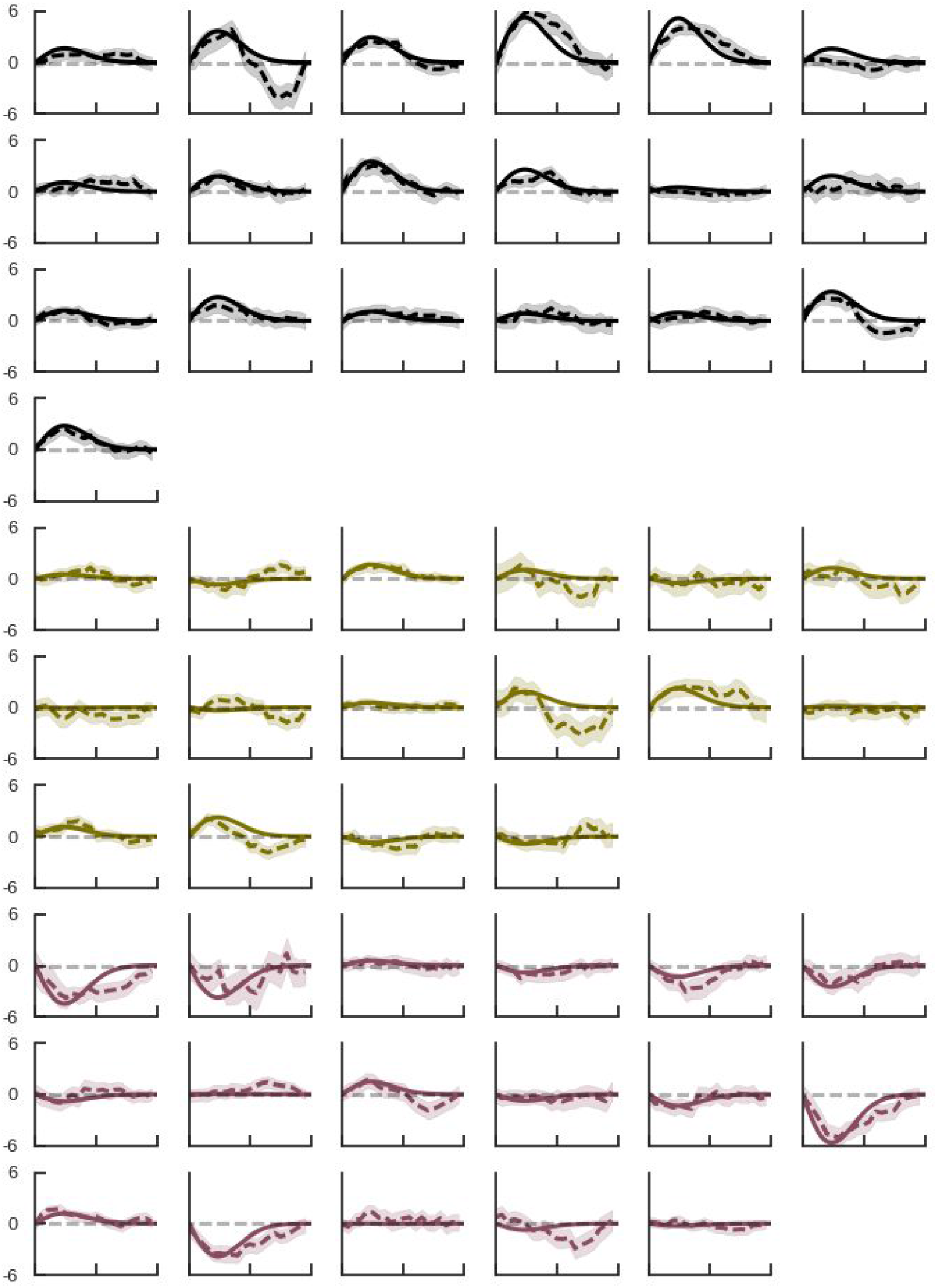
3 sec delay single subject bias and linear mixed model fit. Serial dependence is calculated for each subject as the ‘folded’ error *θ*^e^’ (in degrees, y-axis) for different previous-current distances *θ*^d^ (x-axis, spanning absolute values of 0°-180°) (dashed line; *Methods*). Shading, ± s.e.m. Solid lines show linear model fits (*Methods*, Eq. 1), omitting intercepts and negative values of *θ*^d^ for visualization. Black curves (row 1-4), *ctrl*, green curves (row 5-7), *enc*, purple curves (8-10), *schz*.

**Supplementary Fig. 5.**
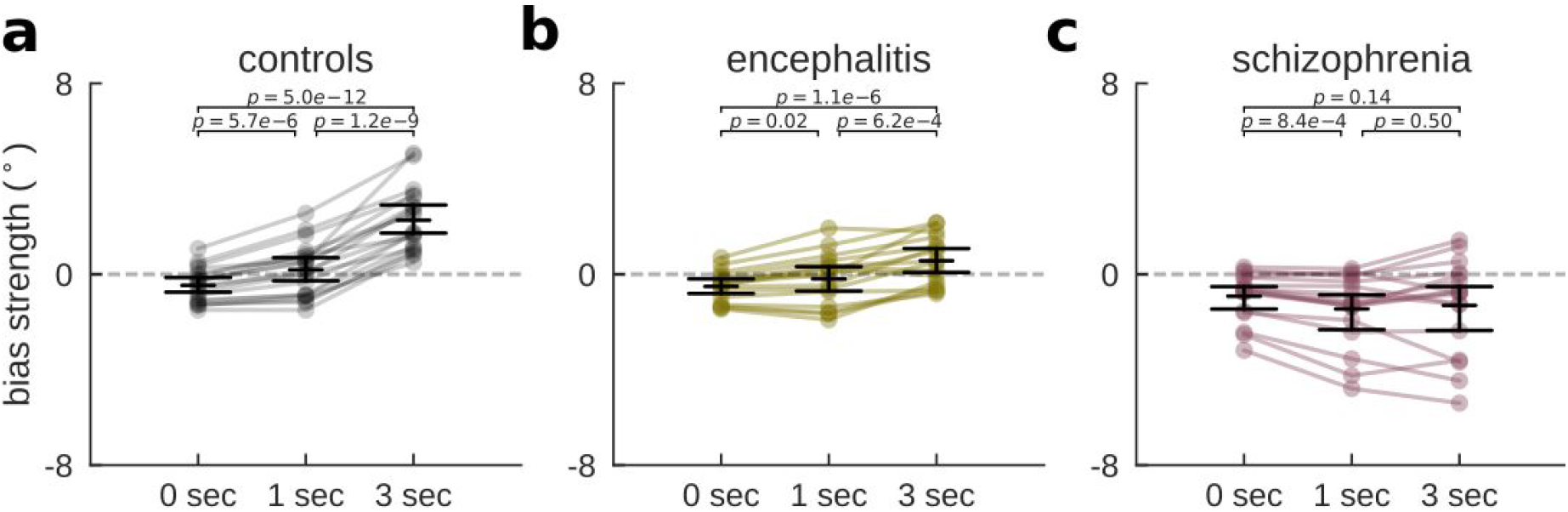
Serial dependence develops as a function of delay length. Individual (random coefficients; dots) and delay-specific group estimates (fixed effects; black horizontal lines indicate mean and bootstrapped 95% C.I. of the mean) of serial dependence. p-values report pairwise comparisons of random coefficients using paired t-tests for n = 19 (ctrl), n = 16 (enc), and n = 17 (schz) patients. **a,** Initially repulsive biases became gradually more attractive with delay length for healthy controls (*Methods*, Eq. 3; *delay* × *DoG(θ*^*d*^), F(2,16.9) = 26.91, p = 5.5e−6; 0 vs 1 sec: t = −6.34, p = 5.7e−6; 1 vs 3 sec: t = −11.34, p = 1.2e−9; 0 vs 3 sec: t = −15.87, p = 5.0e−12) and **b,** for encephalitis patients (*delay × DoG(θ*^*d*^), F(2,22.8) = 5.06, p = 0.015; 0 vs 1 sec: t = −2.71, p = 0.02; 1 vs 3 sec: t = −4.31, p = 6.2e−4; 0 vs 3 sec: t = −7.82, p = 1.1e−6). **c,** schizophrenia patients’ biases did not develop over the course of the delay (*delay × DoG(θ*^*d*^), F(2,18.1) = 1.61, p = 0.23; 0 vs 1 sec: t = 4.10, p = 8.4e-4; 1 vs 3 sec: t = −0.69, p = 0.5; 0 vs 3 sec: t = 1.54, p = 0.14), but stayed repulsive throughout all delay lengths (*DoG(θ*^*d*^), F(1,16.2) = 9.06, p = 0.008).

**Supplementary Fig. 6.**
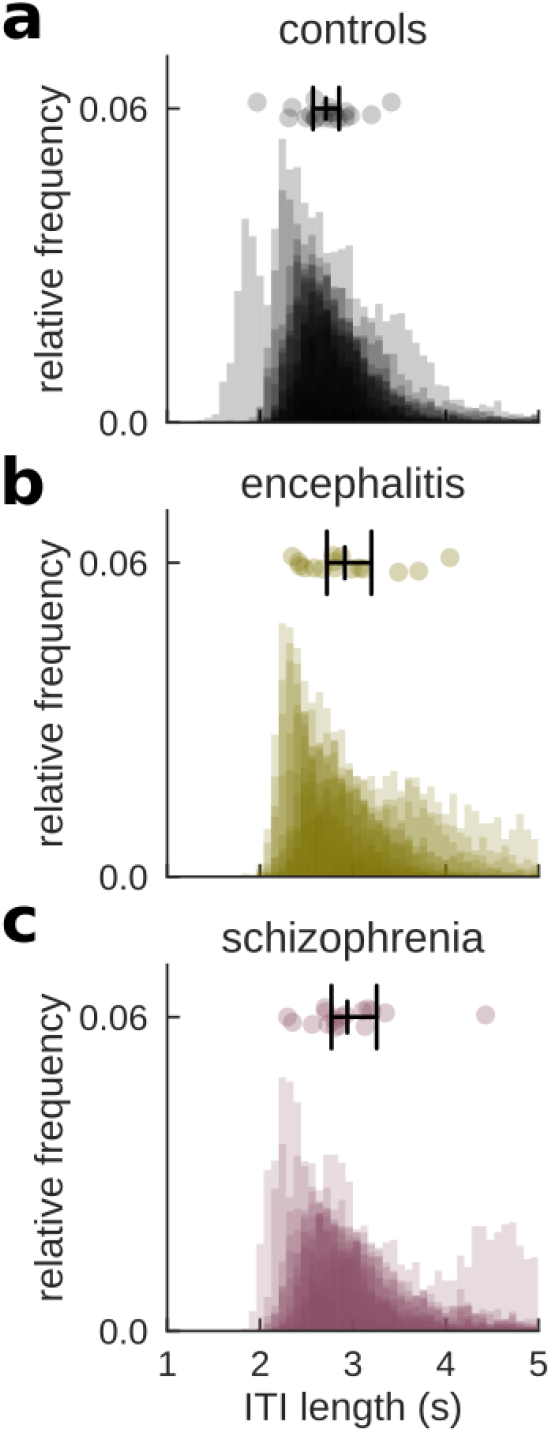
Reduced serial dependence is not explained by group differences in ITI. Histograms of ITI lengths for **a,** control participants **b,** anti-NMDAR encephalitis, and **c,** schizophrenia patients. Each plot shows normalized histograms, transparently overlayed for each participant. Points on top show median ITI lengths for each participant, together with group averages and bootstrapped 95% C.I. (black middle line and error bars). There was a trend for longer median ITIs in patient groups (Kruskal-Wallis test for median ITI length, H = 5.17, p = 0.08; *ctrl*, 2.71±0.33 s; *enc*, 2.91±0.49 s; and *schz*, 3.03±0.46 s; mean±std). Including *ITI* × *DoG(θ*^d^) in our linear model (*Methods*, Eq. 4; ΔAIC = −14.1) did not change group or delay effects of serial dependence (*delay* × *DoG(θ*^d^), F(2,58) = 14.05, p = 1.0e-5; *group* × *DoG(θ*^d^), F(2,50) = 8.10, p = 0.001; *group* × *delay* × *DoG(θ*^d^), F(4,58) = 8.50, p = 1.8e-5), but rather explained additional variance (*ITI × DoG(θ*^*d*^), F(1,7515) = 16.07, p = 6.2e−5).

**Supplementary Fig. 7.**
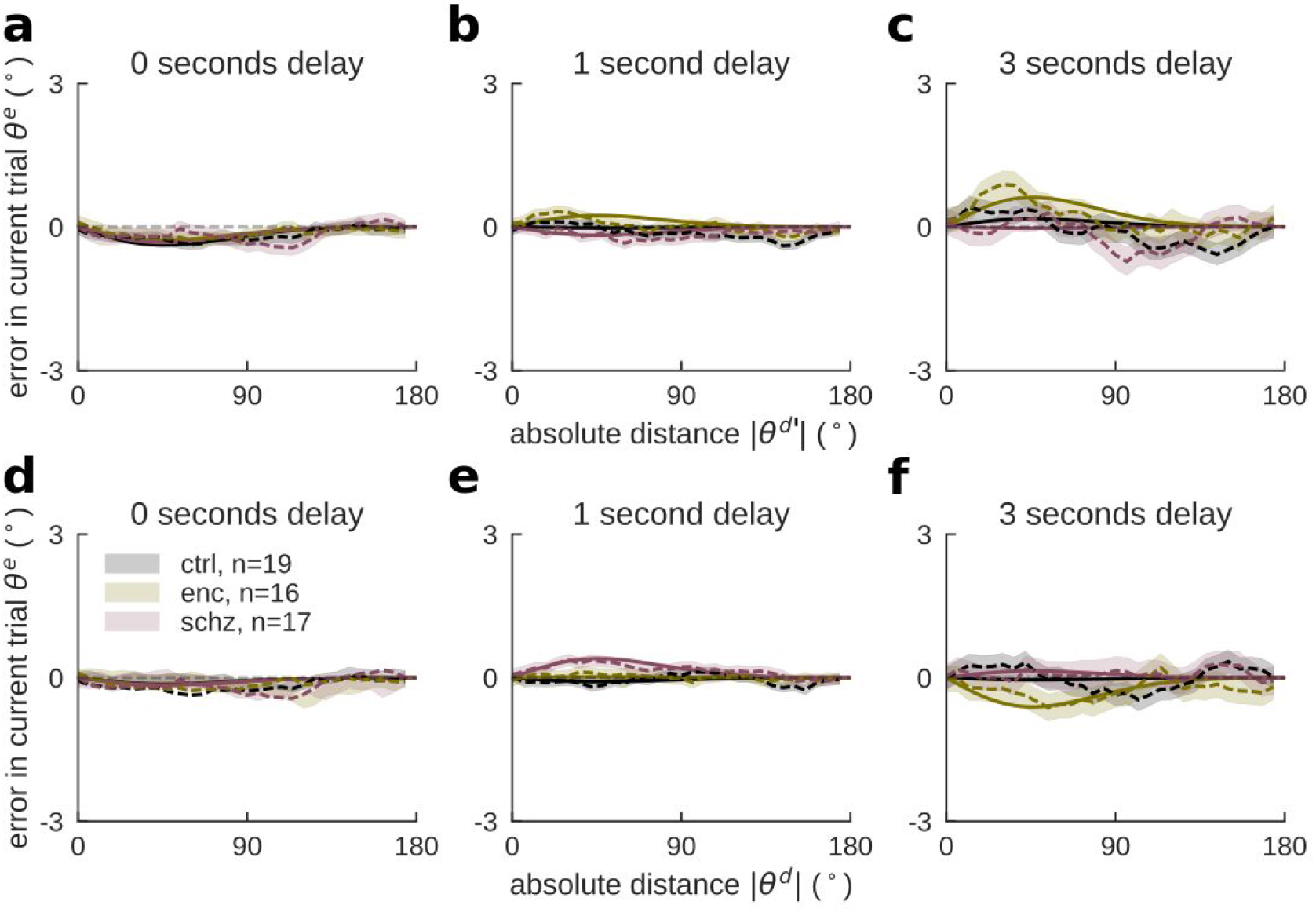
Serial dependence to stimulus n-2 and stimulus n+1. **a,b,c,** An extended linear model with bias terms to both n-1 and n-2 stimuli (adding the
**θ*^d^’*-dependent term *group* × *delay* × *DoG(θ*^*d*^’), *Methods*, Eq. 6; ΔAIC = −4.25) showed significant delay-dependent bias towards the penultimate stimulus (*delay* × *DoG(θ*^*d*^’), F(2,52262) = 5.45, p=0.004). Group differences could not be discarded (*group* × *DoG(θ*^*d*^’), F(2,52269) = 2.87, p = 0.06), but there was no evidence for delay-dependent group differences (*group* × *delay* × *DoG(θ*^*d*^’), F(4,52261) = 0.47, p = 0.76). Groupwise models for each delay showed **a,** significant repulsive bias (*DoG(θ*^*d*^’), F(1,8601.4) = 15.42, p = 8.7e−5) but no group differences for delays of 0 s, (*group* × *DoG(θ*^*d*^’), F(2,8601.5) = 0.10, p = 0.91). **b,** In contrast, groups differed for 1 s delays in absence of overall bias (*DoG(θ*^*d*^’), F(1,34932) = 0.05, p = 0.83; *group* × *DoG(θ*^*d*^’), F(2,34932) = 3.45, p = 0.03), but **c,** not for 3 s delays (*DoG(θ*^*d*^’), F(1,8684) = 3.06, p = 0.08; *group* × *DoG(θ*^*d*^’), F(2,8683.1) = 1.46, p = 0.23). **c, d, e,** We investigated whether serial dependence to stimulus n-1 and group differences in biases could be explained by general response correlations by replacing previous-current distances in Eq. 1 with future-current distances. There was no significant overall bias towards future stimuli (*DoG(θ*^*d*^), F(1,80) = 0.74, p = 0.39; *delay* × *DoG(θ*^*d*^), F(2,119) = 2.40, p = 0.09; *group* × *DoG(θ*^*d*^), F(2,80) = 1.80, p = 0.17; *group* × *delay* × *DoG(θ*^*d*^), F(4,119) = 1.05, p = 0.38), indicating non-significant contributions of general response correlations between trials to the reported group and delay effects of serial dependence.

**Supplementary Fig. 8.**
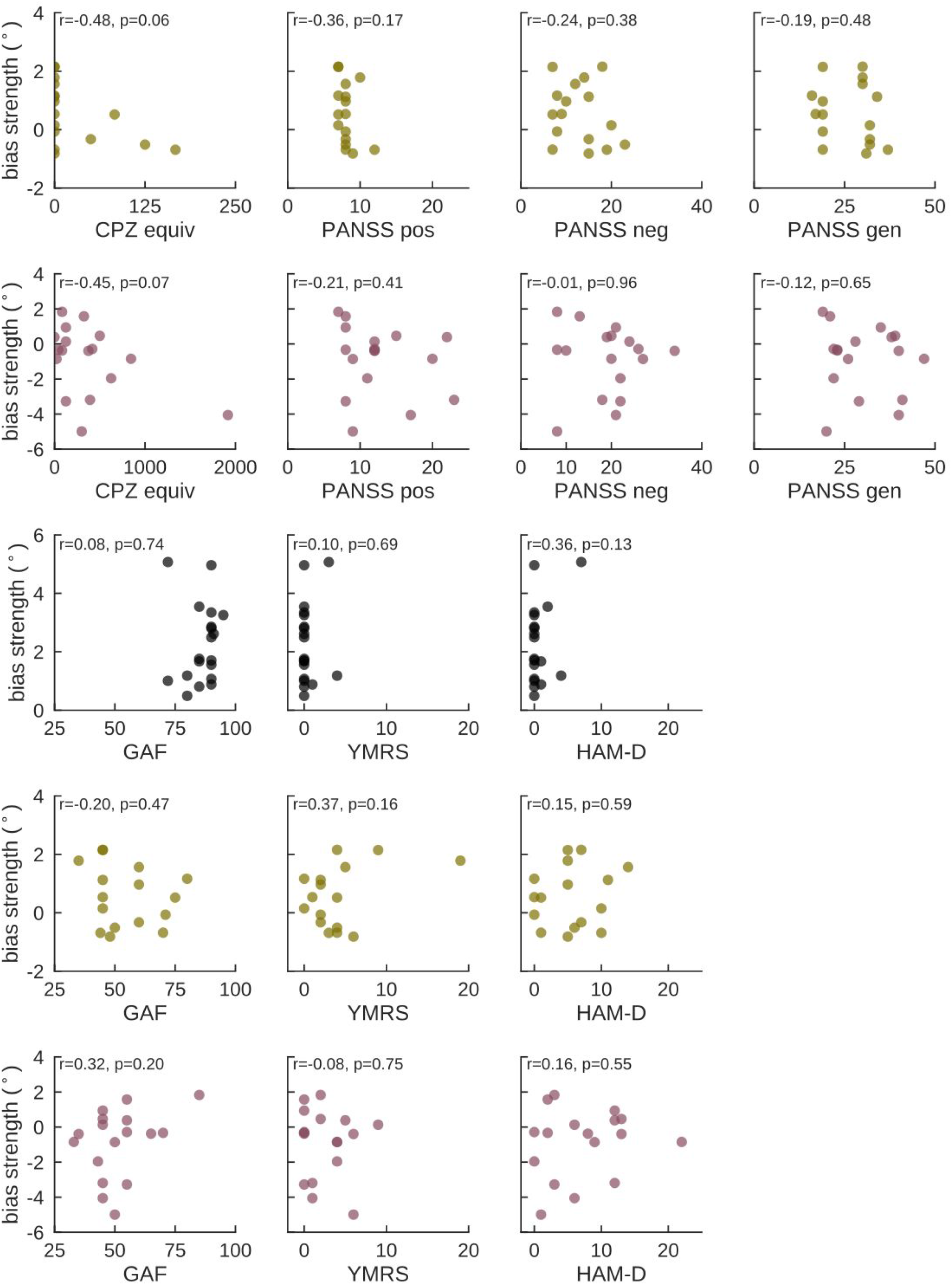
Correlations of serial dependence in 3 s delay trials with clinical scales. For each group, we correlated individual bias coefficients for 3 s delay trials (random effects) with clinical measures (*Methods* for description of administered tests). The strength of serial dependence did not correlate significantly with clinical scales. Correlations were calculated using Pearson’s r for n = 19 (ctrl, black, row 3), n = 16 (enc, green, rows 1 and 4), and n = 17 (schz, purple, rows 2 and 5). Correlations with antipsychotic medication (CPZ equiv) reached marginal significance for both encephalitis and schizophrenia. To test whether medication could account for group differences in delay-dependent bias, we included a transversal estimate of antipsychotic medication, *CPZ*, as a covariate in our linear model (*Methods*, Eq. 5; ΔAIC = −2.7). Antipsychotic medication explained a significant amount of variance in delay-dependent bias (*CPZ* × *delay* × *DoG(θ*^*d*^), F(3,60) = 3.07, p = .03), but did not change the pattern of results (*delay* × *DoG(θ*^*d*^), F(2,62) = 17.58, p = 8.9e−7; *group* × *DoG(θ*^*d*^), F(2,48) = 3.92, p = 0.03; *group* × *delay* × *DoG(θ*^*d*^), F(4,62) = 4.45, p = 0.003). Measures: GAF (Global Assessment of Functioning Scale ^45^), YMRS (Young Mania Rating Scale ^43^), HAM-D (Hamilton Depression Rating Scale ^44^), PANSS (Positive and Negative Syndrome Scale ^42^) Positive, Negative and General Psychopathology Scale, CPZ equiv (transversal estimate of antipsychotic medication as chlorpromazine equivalent).

**Supplementary Fig. 9.**
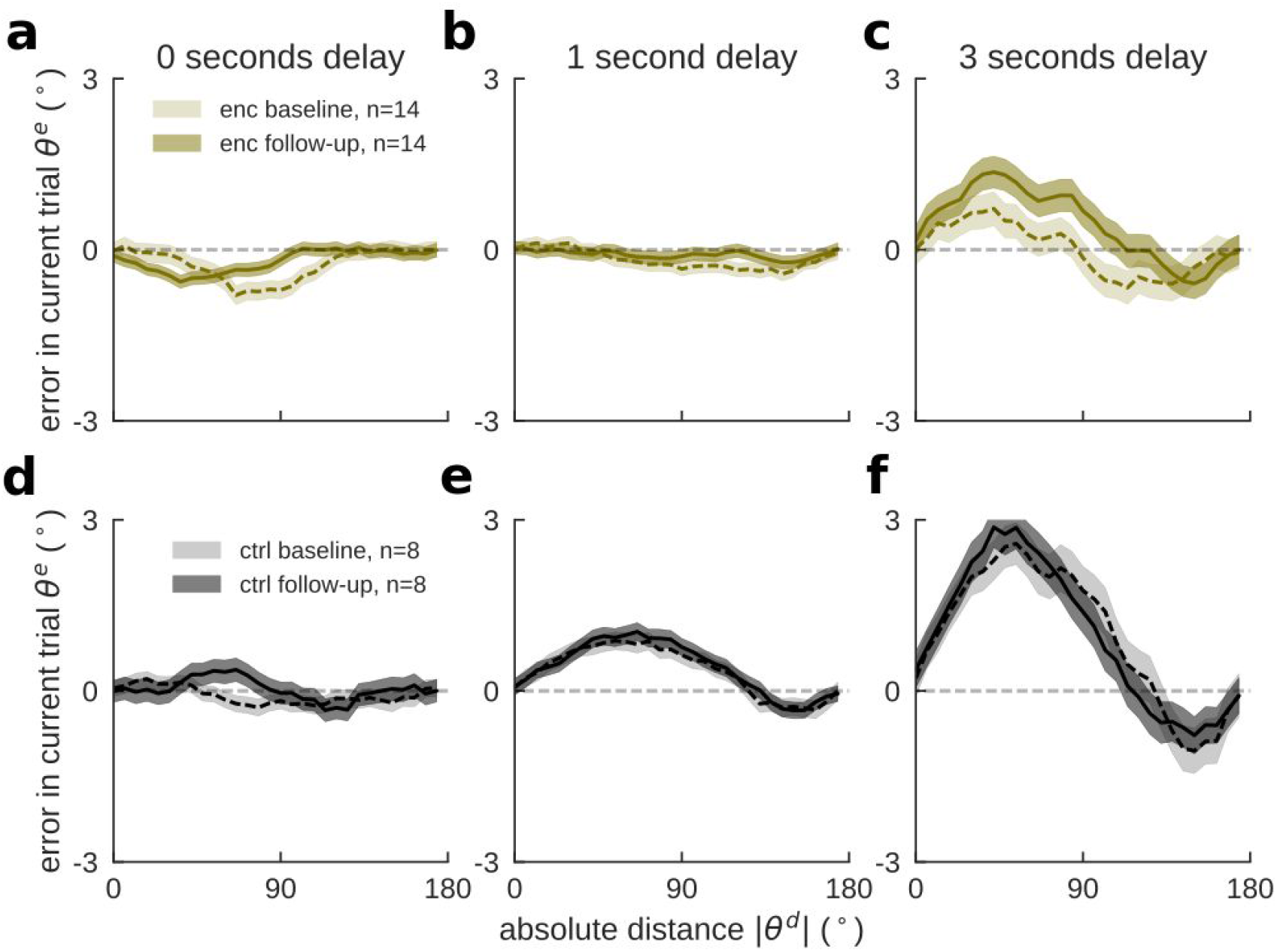
Serial dependence increases with encephalitis patients’ recovery. We performed a comparison of baseline and follow-up sessions (*Methods*, Eq. 7) for n=14 encephalitis patients (enc, Supplementary Table 2) and n=8 controls (ctrl). Although the four-way interaction did not reach significance (*session* × *group* × *delay* × *DoG(θ*^*d*^), F(2,30124) = 0.79, p = 0.45), group-wise models showed a normalization of biases in encephalitis patients’ (**a,b,c,** *session* × *delay* × *DoG(θ*^*d*^), F(2,30124) = 3.07, p = 0.046), and not in healthy controls (**d,e,f,** F(2,16311) = 0.10, p = 0.90). A delay-wise comparison of encephalitis patients’ baseline and follow-up values showed that this difference was driven by biases in 3 second delays (*session × DoG(*θ*^d^)*, F(1,5030) = 4.43, p = 0.035), while biases in 0 and 1 second delays did not change (F(1,5030) = 0.15, p = 0.69, and F(1,20064) = 0.05, p = 0.81, respectively). Note that due to the increased complexity of the model and the limited sample size, we could not estimate random effects in this comparison.

**Supplementary Fig. 10.**
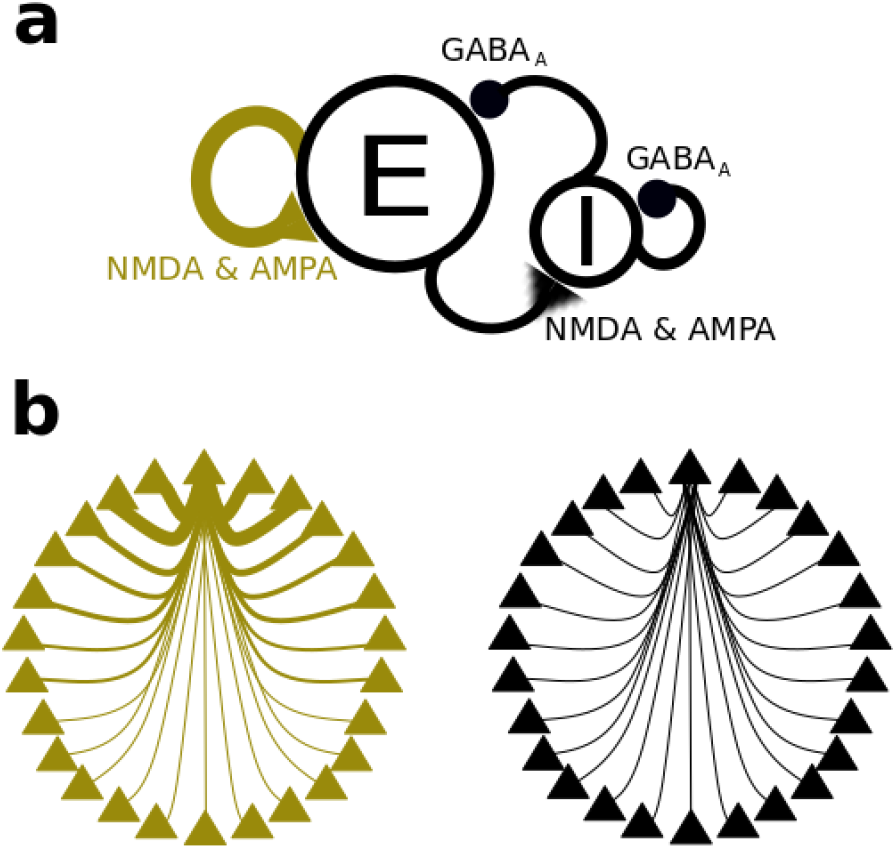
Network scheme and connectivity profile. **a,** Scheme of spiking neural network, consisting of 1024 excitatory and 256 inhibitory neurons. Neurons from both pools were connected in an all-to-all fashion, with excitatory connections governed by NMDA and AMPA dynamics, and inhibitory connections governed by GABAA dynamics. STP affected recurrent excitatory connections. **b,** Weight profiles for recurrent excitatory (green) and all other connections (black). For recurrent connections, weights between neurons preferring similar locations were higher, while more distant neurons were only weakly connected. All other connections had flat connectivity profiles, with equal weights between similar and dissimilar neurons.

**Supplementary Fig. 11.**
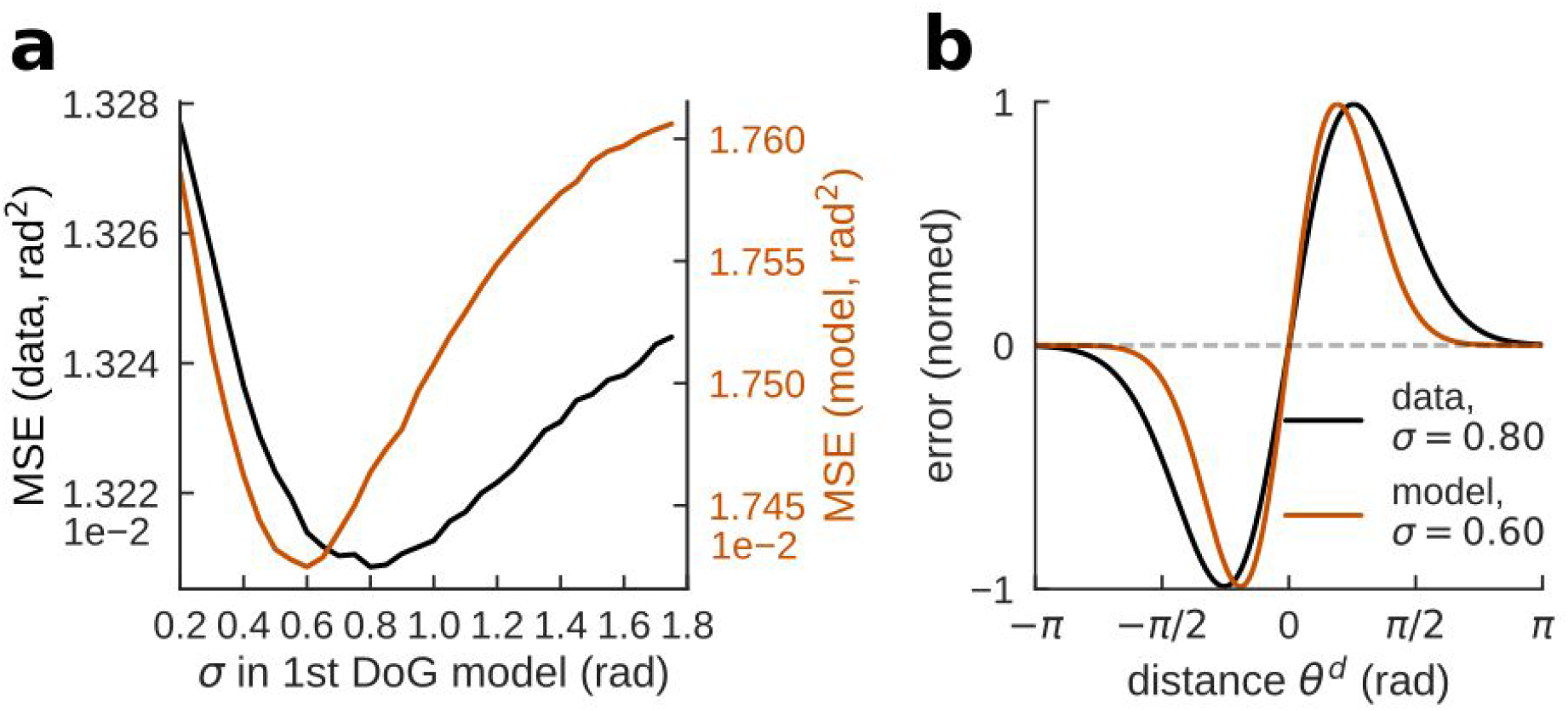
Hyperparameter cross-validation and model selection. **a,** Mean squared error for stratified hyperparameter optimization using cross-validation (1000 repetitions, training set size = .33 from each subject) for data (black) and for the network model model (orange; cross-validation with 1000 repetitions, training set size = .33 of 21.000 simulated trials with baseline STP and conductance parameters, corresponding to the control condition in Fig. 3). Hyperparameters are different values of variance *σ* (in radians) of the underlying Gaussian with location hyperparameter *μ* = 0. **b,** Shape of first-derivative-of-Gaussian fits with optimal hyperparameter *σ* and *μ* = 0 for data (black) and model (orange). Hyperparameter cross-validation for neural network simulations was carried out for the default parameters of STP, g_EI_ g_EE_ as reported in *Methods*. Note that in b, signed previous-current distances are indicated in radians as used in the linear model.

**Supplementary Fig. 12.**
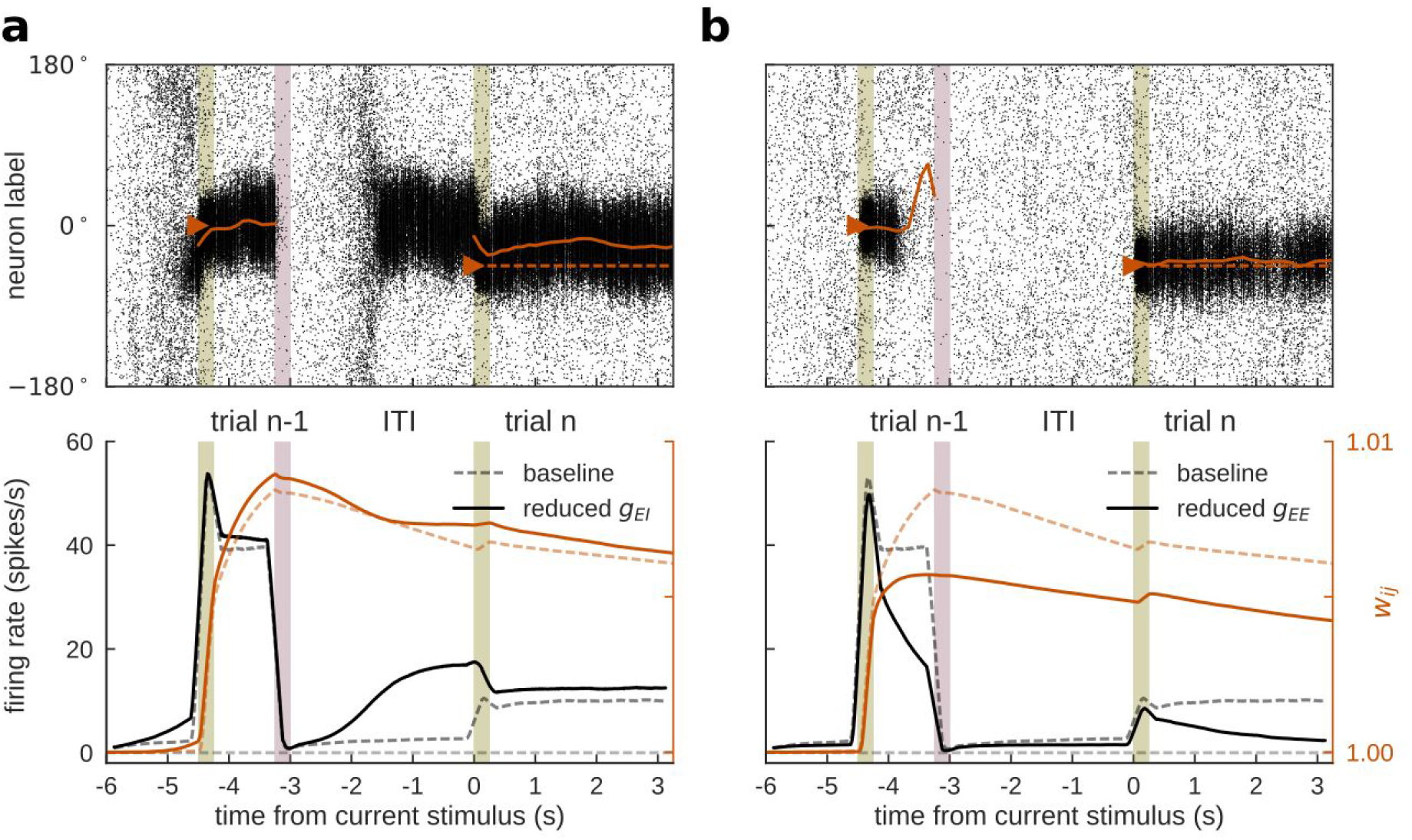
Network behavior for reduced g_EI_ and g_EE_. **a,** For reduced g_EI_, network activity was disinhibited and baseline firing became unstable. Spontaneous activity bumps emerged in the ITI (upper panel), often in neurons that had been active during the previous delay. Lower panel shows firing rates and STP traces at neurons selective to stimuli appearing at 0° for the baseline condition (0% reduction, dashed lines) and the disinhibited condition (1% reduction, solid lines), averaged over 1000 trials in which the second stimulus appeared at randomized locations. **b,** For reduced g_EE_, delay firing became unstable and active working memory representations were lost over the delay (upper panel). Lower panel analogous to **a,** for the baseline condition (0% reduction, dashed lines) and the condition of reduced excitation (1% reduction, solid lines). Lower panels were computed as in Fig. 2 but including trials for which the same neurons were coactive in the two successive trials. This explains the difference between dashed lines here and in Fig. 2b, trial *n*.

## Supplementary Tables

**Supplementary Table 1.**
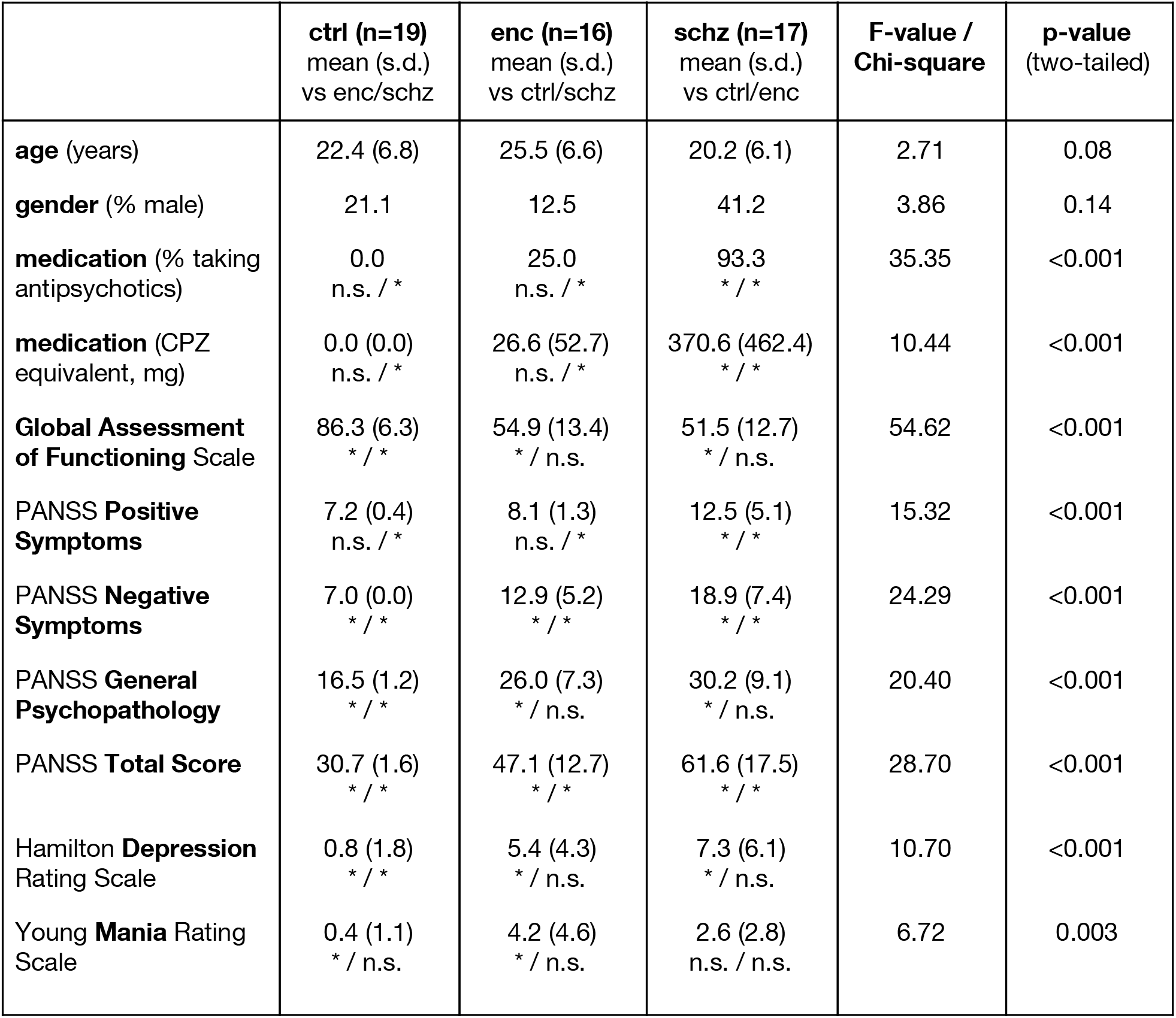
Clinical and demographic statistics of the population. Measures: GAF (Global Assessment of Functioning Scale ^45^), YMRS (Young Mania Rating Scale ^43^), HAM-D (Hamilton Depression Rating Scale ^44^), PANSS (Positive and Negative Syndrome Scale ^42^) Positive, Negative and General Psychopathology Scale, CPZ equiv (transversal estimate of antipsychotic medication as chlorpromazine equivalent). The significance of pairwise post-hoc Tukey/Bonferroni-corrected chi-square tests is reported below group mean and s.d. (“n.s.” marking non-significant comparisons, and “ * ” significant comparisons with FWE=0.05).

**Supplementary Table 2.** Baseline/follow-up comparison of anti-NMDAR encephalitis patients. Measures: GAF (Global Assessment of Functioning Scale ^45^), YMRS (Young Mania Rating Scale ^43^), HAM-D (Hamilton Depression Rating Scale ^44^), PANSS (Positive and Negative Syndrome Scale ^42^) Positive, Negative and General Psychopathology Scale, CPZ equiv (transversal estimate of antipsychotic medication as chlorpromazine equivalent).

|  | **baseline (n=14)**<br>mean (s.d.) | **follow-up (n=14)**<br>mean (s.d.) | **t-value /**<br>**Chi-square** | **p-value**<br>(two-tailed) |
| --- | --- | --- | --- | --- |
| **medication** (% taking antipsychotics) | 21.4 | 7.1 | 0.29 | 0.59 |
| **medication** (CPZ equivalent, mg) | 21.4 (48.7) | 3.0 (11.2) | 1.34 | 0.20 |
| **Global Assessment of Functioning Scale** | 53.4 (12.4) | 72.1 (13.7) | -4.69*** | <0.001 |
| **PANSS Positive Symptoms** | 8.2 (1.4) | 8.2 (0.9) | 0.00 | 1.00 |
| **PANSS Negative Symptoms** | 12.6 (4.6) | 8.6 (2.4) | 3.24** | 0.006 |
| **PANSS General Psychopathology** | 26.3 (7.1) | 20.6 (3.2) | 2.86* | 0.01 |
| **PANSS Total Score** | 47.1 (12.1) | 37.4 (5.5) | 2.85* | 0.01 |
| **Hamilton Depression Rating Scale** | 5.8 (4.4) | 3.4 (4.0) | 2.12 | 0.05 |
| **Young Mania Rating Scale** | 4.5 (4.8) | 3.1 (2.3) | 1.11 | 0.29 |

